# With No Lysine Kinase 1 Promotes Right Ventricular Dysfunction Via Glucotoxicity

**DOI:** 10.1101/2021.06.22.449476

**Authors:** Sasha Z. Prisco, Megan Eklund, Thenappan Thenappan, Kurt W. Prins

**Author notes:** **Author for Correspondence** Kurt Prins, MD, PhD, Assistant Professor of Medicine, Lillehei Heart Institute, Cardiovascular Division, University of Minnesota Medical School, Minneapolis, MN 5545. Funding: SZP is funded by NIH F32 HL154533, NIH T32 HL144472, a University of Minnesota Clinical and Translational Science award (NIH UL1 TR002494), and a University of Minnesota Medical School Academic Investment Educational Program Grant; TT is funded by the Cardiovascular Medical Research and Education Fund and the University of Minnesota Futures Grant; KWP is funded by NIH K08 HL140100, the Cardiovascular Medical Research and Education Fund, a Lillehei Heart Institute Cardiovascular Seed Grant, the University of Minnesota Faculty Research Development Grant, the United Therapeutics Jenesis Award, and an American Lung Association Innovative Award IA-816386. The content is solely the responsibility of the authors and does not represent the official views of the NIH or any other funding sources.

## Abstract

**Objectives:** Investigate how WNK1 inhibition modulates glucotoxicity, mitochondrial/peroxisomal protein regulation and metabolism, and right ventricular (RV) function in pulmonary arterial hypertension (PAH). Determine how hypochloremia impacts RV function in PAH patients.

**Background:** In PAH-induced RV failure, GLUT1/GLUT4 expression is elevated, which increases glucose uptake and glycolytic flux to compensate for mitochondrial dysfunction. However, the resultant consequences of the glucose-mediated post-translational modifications (PTM), protein O-GlcNAcylation/glycation in RV failure are understudied. WNK1, a chloride-sensitive kinase, increases GLUT1/GLUT4 expression in skeletal muscle, but its regulation in RV dysfunction is unexplored.

**Methods:** Rats were treated with WNK463 (small molecule WNK inhibitor) or vehicle starting two weeks after monocrotaline injection. Immunoblots quantified protein abundance/PTMs. Mitochondrial/peroxisomal proteomics and global metabolomics evaluated glucose metabolism and mitochondrial/peroxisomal function. Pulmonary vascular and cardiac histology, echocardiography, and pressure-volume loop analysis quantified RV function and PAH severity. Finally, the relationship between hypochloremia, a WNK1-activating state, and RV function was evaluated in 217 PAH patients.

**Results:** WNK463 decreased WNK1/GLUT1/GLUT4 expression, normalized glucose metabolite levels, which dampened excess protein O-GlcNAcylation/glycation. Integration of RV mitochondrial/peroxisomal proteomics and metabolomics identified fatty acid oxidation (FAO) as the most dysregulated metabolic pathway. WNK463 enhanced FAO as demonstrated by increased expression of mitochondrial FAO proteins and normalization of RV acylcarnitines. WNK463 reduced glutaminolysis induction and lipotoxicity, two secondary consequences of diminished FAO. WNK463 augmented RV systolic and diastolic function independent of pulmonary vascular disease severity. In PAH patients, hypochloremia resulted in more severe RV dysfunction.

**Conclusions:** WNK463 combated RV glucotoxicity and impaired FAO, which directly improved RV function.

**Highlights:**

- Small molecule inhibition of WNK1 (WNK463) signaling mitigates upregulation of the membrane glucose channels GLUT1 and GLUT4, restores levels of several glucose metabolites, and normalizes protein O-GlcNAcylation and glycation in the RV.
- Quantitative proteomics of RV mitochondrial enrichments shows WNK463 treatment prevents downregulation of mitochondrial enzymes in the tricarboxylic acid cycle, fatty acid oxidation pathway, and the electron transport chain complexes.
- Integration of proteomics and metabolomics analysis reveals WNK463 reduces glutaminolysis induction and lipotoxicity due to impaired fatty acid oxidation
- WNK463 augments RV systolic and diastolic function independent of PAH severity.
- Hypochloremia, a condition of predicted WNK1 activation, in PAH patients results in more severe RV dysfunction.

## Introduction

Pulmonary arterial hypertension (PAH) is a proliferative vasculopathy of the resistance pulmonary arteries that leads to elevated pulmonary arterial pressures (1). The pathological changes in the pulmonary vascular bed augment the workload of the right ventricle (RV), which ultimately results in RV dysfunction (RVD) (2,3). Metabolic derangements are the most robustly characterized phenotypes of RVD in PAH (4,5) with resultant increases in RV glucose uptake in both preclinical (6,7) and human PAH (8,9). Interestingly, RV function is inversely associated with RV glucose uptake (8), which implies excess intracellular glucose may have deleterious effects. While glucose metabolism primarily generates adenosine triphosphate (ATP), the hexosamine biosynthetic pathway converts glucose to UDP-*N*-acetylglucosamine (UDP-GlcNAc) (10). UDP-GlcNAc is used to enzymatically post-translationally modify (PTM) serine or tyrosine residues, a process known as protein O-GlcNAcylation (10). Moreover, 1-2% of glucose metabolized through glycolysis results in formation of methylglyoxal, a highly reactive dicarbonyl that can nonenzymatically modify proteins (protein glycation) (11). Interestingly, both excess O-GlcNAcylation and glycation promote left ventricular (LV) cardiomyocyte mitochondrial dysfunction (12,13), but the role of these PTMs in RVD is relatively unexplored.

Two clinical studies show hypochloremia identifies high-risk PAH patients, which may have direct relevance to glucose metabolism. Naal *et al*. showed that hypochloremia is independently associated with increased mortality (14). We also demonstrated that hypochloremia is independently associated with increased mortality and measures of RV failure in a multi-center study (15). The <u>W</u>ith-<u>N</u>o-<u>L</u>ysine (WNK) kinase proteins are a family of signaling kinases activated in the setting of low intracellular chloride levels (16,17). In the heart, WNK1 is the predominant isoform expressed (17), but studies of WNK1 in cardiac diseases are lacking. WNK1 function is better understood in skeletal muscle as previous studies show WNK1 promotes membrane localization of the glucose channels, GLUT1 (18) and GLUT4 (19) via activation of 160 kDa substrate of the Akt Ser/Thr kinase (AS160), a RAB GTPase-activating protein. Thus, these data suggest hypochloremia could result in WNK1 activation and subsequently modulate RV glucose metabolism.

Here, we investigated the effects of WNK inhibition on RV glucotoxicity, metabolism, and function in preclinical PAH. We implemented a translational approach by treating monocrotaline (MCT) rats two weeks after the development of PAH and comprehensively examined the RV’s response to WNK inhibition using immunoblots, proteomics, metabolomics, echocardiography, and pressure-volume (PV) loop analysis. Finally, we assessed how hypochloremia, a WNK1-activating state, modulated RV function in 217 PAH patients.

## Methods

Extended methods are in the **Supplemental Materials**. In brief, male Sprague Dawley rats received a single subcutaneous injection of MCT. Two weeks after MCT injection, rats were given daily intraperitoneal injections of either 3 mg/kg WNK463 or vehicle. Immunoblots of RV extracts were completed as previously described (20). Antibodies used in this study are listed in **Supplemental Table 1**. Complete Western blot images are shown in **Supplemental Figure 1**.

RV mitochondrial/peroxisomal enrichment fractions were used for quantitative mass spectrometry as described in the **Supplemental Methods**. Metabolomic profiling of frozen RV tissues was completed by Metabolon, Inc. (Durham, NC).

Lung and cardiac histology, echocardiography (**Supplemental Figure 2**), and PV loops (**Supplemental Figure 3**) quantified the effects of WNK463 on RV structure/function and pulmonary vascular disease severity.

Finally, we determined how hypochloremia impacted RV function in a PAH cohort of 217 patients (**Supplemental Table 2**).

## Results

### WNK Inhibition Mitigated GLUT1 and GLUT4 Upregulation and Reduced Levels of Multiple RV Glucose Metabolites

Immunoblots showed MCT-Vehicle (MCT-V) RVs had elevated levels of WNK1, GLUT1, GLUT4, phosphorylated (active form) of AS160, and the ratio of pAS160/AS160, which was mitigated by WNK463 (**Figure 1A**). Confocal microscopy revealed increased WNK1 immunoreactivity (**Figure 1B**) and membrane localization of GLUT1 (**Figure 1C**) and GLUT4 (**Figure 1D**) in MCT-V RVs. Importantly, WNK463 reduced WNK1 staining intensity (**Figure 1B**) and GLUT1 (**Figure 1C**) and GLUT4 membrane enrichment (**Figure 1D**). Metabolomics profiling quantified the effects of WNK inhibition on glucose metabolites in the hexosamine biosynthetic, glycolytic, and pentose phosphate pathways (**Figure 1E**). In MCT-V RVs, the levels of the end-products in the hexosamine biosynthetic (UDP-GlcNAc) (**Figure 1F**), glycolytic (pyruvate) (**Figure 1G**), and pentose phosphate pathways (erythrose 4-phosphate) (**Figure 1H**) were all higher than controls, but WNK463 restored the concentration of these metabolites (**Figure 1F-G**). Thus, these data support a role of WNK1 in regulating RV glucose handling and metabolism.

**Figure 1:**
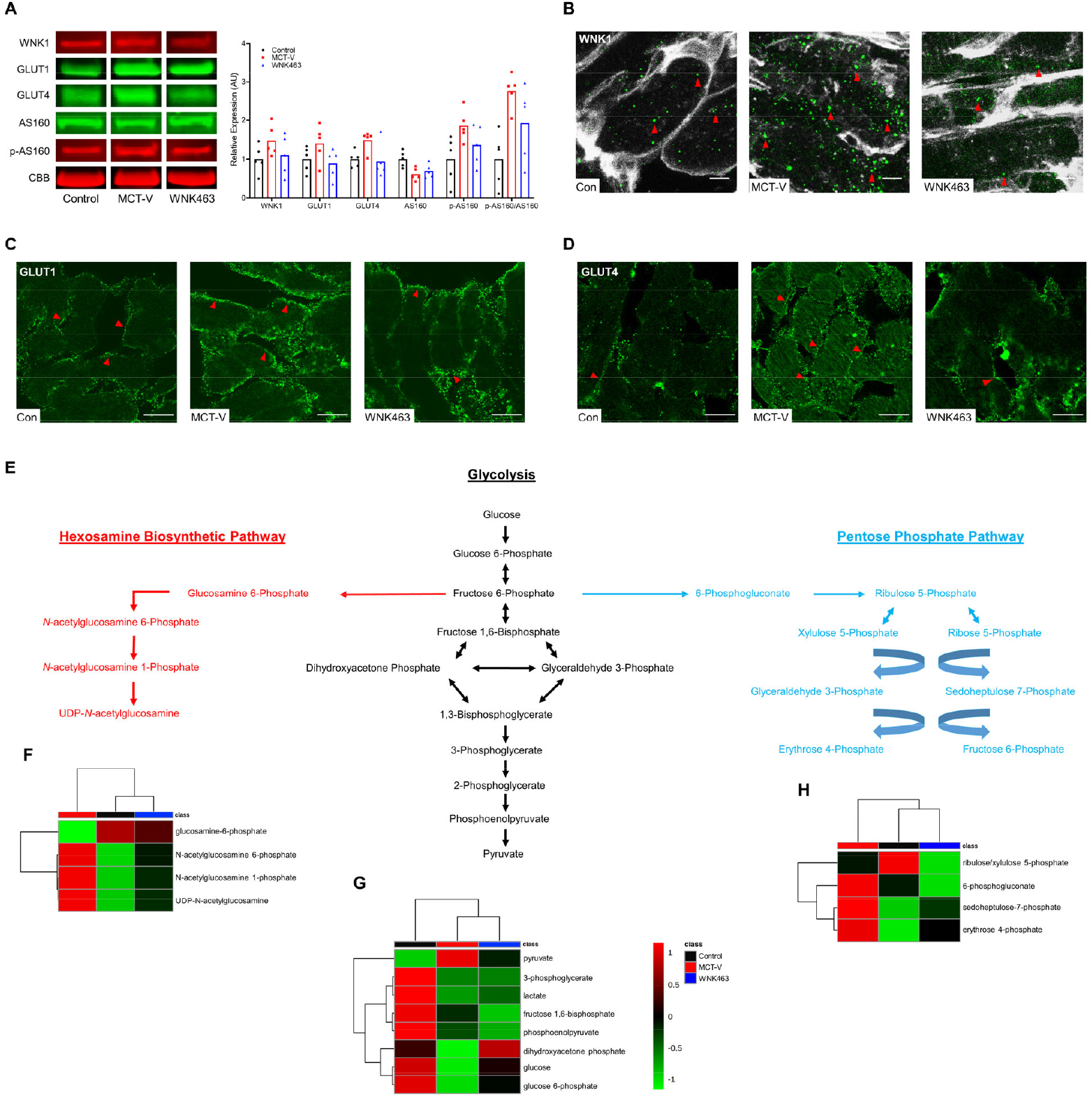
WNK Inhibition Combats RV Glucotoxicity in MCT Rats. (**A**) Representative Western blots and quantification of protein abundance in RV extracts from control, MCT-V, and WNK463 rats demonstrates WNK463 partially normalizes expression of WNK1 (Control: 1.0±0.4, MCT-V: 1.5±0.4, WNK463: 1.1±0.5 expression relative to control, *n*=5 RVs per group for all proteins evaluated), GLUT1 (Control: 1.0±0.3, MCT-V: 1.4±0.5, WNK463: 0.9±0.4), GLUT4 (Control: 1.0±0.2, MCT-V: 1.5±0.3, WNK463: 0.9±0.4), AS160 (Control: 1.0±0.2, MCT-V: 0.6±0.2, WNK463: 0.7±0.2), phosphorylated AS160 (Control: 1.0±0.6, MCT-V: 1.9±0.4, WNK463: 1.4±0.5), and phosphorylated AS160 normalized to AS160 (Control: 1.0±0.7, MCT-V: 2.8±0.5, WNK463: 1.9±1.0). Data presented as expression relative to control. Representative immunofluorescence images from RV free wall sections show WNK463 reduces the amount of (**B**) cytoplasmic WNK1 expression (red arrows) (wheat germ agglutinin staining in white) and (**C**) GLUT1 receptors (red arrows) and (**D**) GLUT4 receptors (red arrows) at the cell membrane. Scale bar 10 µm in (**B**) and 20 µm in (**C**) and (**D**). (**E**) Hexosamine biosynthetic pathway, glycolysis, and pentose phosphate pathway intermediates are outlined. RV metabolomics studies demonstrate WNK463 restores the levels of (**F**) hexosamine biosynthesis pathway, (**G**) glycolysis, and (**H**) pentose phosphate pathway intermediates as depicted by hierarchical cluster analysis.

### WNK Inhibition Combated Excess Protein O-GlcNAcylation and Glycation

Next, we examined how WNK463 treatment altered protein O-GlcNAcylation/glycation in the RV. O-GlcNAcylation was increased in MCT-V RVs, which was mitigated by WNK463 treatment (**Figure 2A-B**). Furthermore, WNK463 reduced expression of O-GlcNAcase (OGA) and glutamine-fructose-6-phosphate transaminase 1 (GFAT1) (**Figure 2A-B**). Confocal microscopy showed WNK463 decreased intracellular cardiomyocyte O-GlcNAcylation (**Figure 2C**). Next, we investigated the effects of WNK inhibition on protein glycation and the proteins that modulate protein glycation. WNK463 curtailed excess protein glycation that was observed in MCT-V without drastically changing expression of DJ-1, a protein deglycase, glyoxalase 1 (GLO1) and GLO2, proteins that catabolize methylglyoxal, the glucose metabolite responsible for protein glycation (**Figure 2D-E**) (21). Thus, WNK463 decreased protein O-GlcNAcylation/glycation, further suggesting inhibition of RV glucotoxicity.

**Figure 2:**
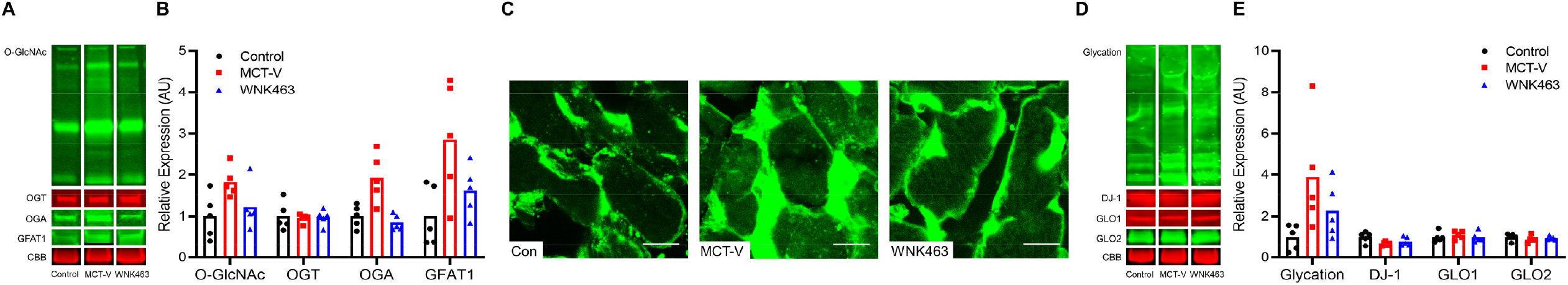
Protein O-GlcNAcylation and Glycation are Blunted by WNK463. (**A**) Representative Western blots and (**B**) quantification of protein abundance in RV extracts from control, MCT-V, and WNK463 rats demonstrate WNK463 normalizes protein O-GlcNAcylation (Control: 1.0±0.5, MCT-V: 1.8±0.4, WNK463: 1.2±0.5 expression relative to control, *n*=5 RVs per group for all proteins assessed), OGA (Control: 1.0±0.3, MCT-V: 1.9±0.6, WNK463: 0.9±0.2), and GFAT1 (Control: 1.0±0.7, MCT-V: 2.9±1.4, WNK463: 1.6±0.6) expression and did not change OGT abundance (Control: 1.0±0.3, MCT-V: 1.0±0.1, WNK463: 1.0±0.2). Data presented as expression relative to control. (**C**) Representative confocal micrographs of RV free wall sections stained with succinylated WGA show increased intracellular O-GlcNAcylation signal in MCT-V, which is reduced by WNK463. Scale bar 10 µm. (**D**) Representative Western blots and (**E**) quantification of protein glycation, DJ-1, GLO1, and GLO2 reveal WNK463 partially normalizes total protein glycation (Control: 1.0±0.6, MCT-V: 3.9±2.7, WNK463: 2.3±1.3) without altering DJ-1, GLO1, and GLO2 abundance in the RV.

### Proteomics Revealed WNK463 Prevented Dysregulation of Proteins in the Tricarboxylic Acid (TCA) Cycle, Fatty Acid Oxidation (FAO), and Electron Transport Chain (ETC) Complexes

Because existing data suggest both excess protein O-GlcNAcylation and glycation negatively regulate mitochondrial function, we performed quantitative proteomic profiling to examine how WNK463 altered mitochondrial/peroxisomal protein homeostasis. We identified 2970 total proteins in our extracts, and 1203 proteins had significant differences in abundance. Principal component analysis revealed WNK463 shifted the proteomic signature towards control (**Figure 3A**). Hierarchical cluster analysis corroborated this finding (**Figure 3B**). Next, we determined how WNK463 altered expression of TCA cycle enzymes. WNK463 increased expression of the enzymes Sdhb (succinate dehydrogenase complex iron sulfur subunit B), Aco2 (aconitase 2), and Sdha (succinate dehydrogenase complex flavoprotein subunit A) when compared to MCT-V (**Figure 3C**). Then, we probed mitochondrial FAO enzyme regulation. MCT-V rats had lower levels of multiple FAO proteins, which WNK463 combated (**Figure 3D**). Finally, the protein subunits of the ETC complexes I-V were evaluated. MCT-V RVs had a reduction in nearly all subunits of complexes I-V, but WNK463 mitigated the downregulation of almost all of these proteins (**Figure 3E-I**). In summary, proteomics analysis revealed WNK463 restored protein abundance of several enzymes in the TCA cycle, FAO pathway, and complexes I-V of the ETC.

**Figure 3:**
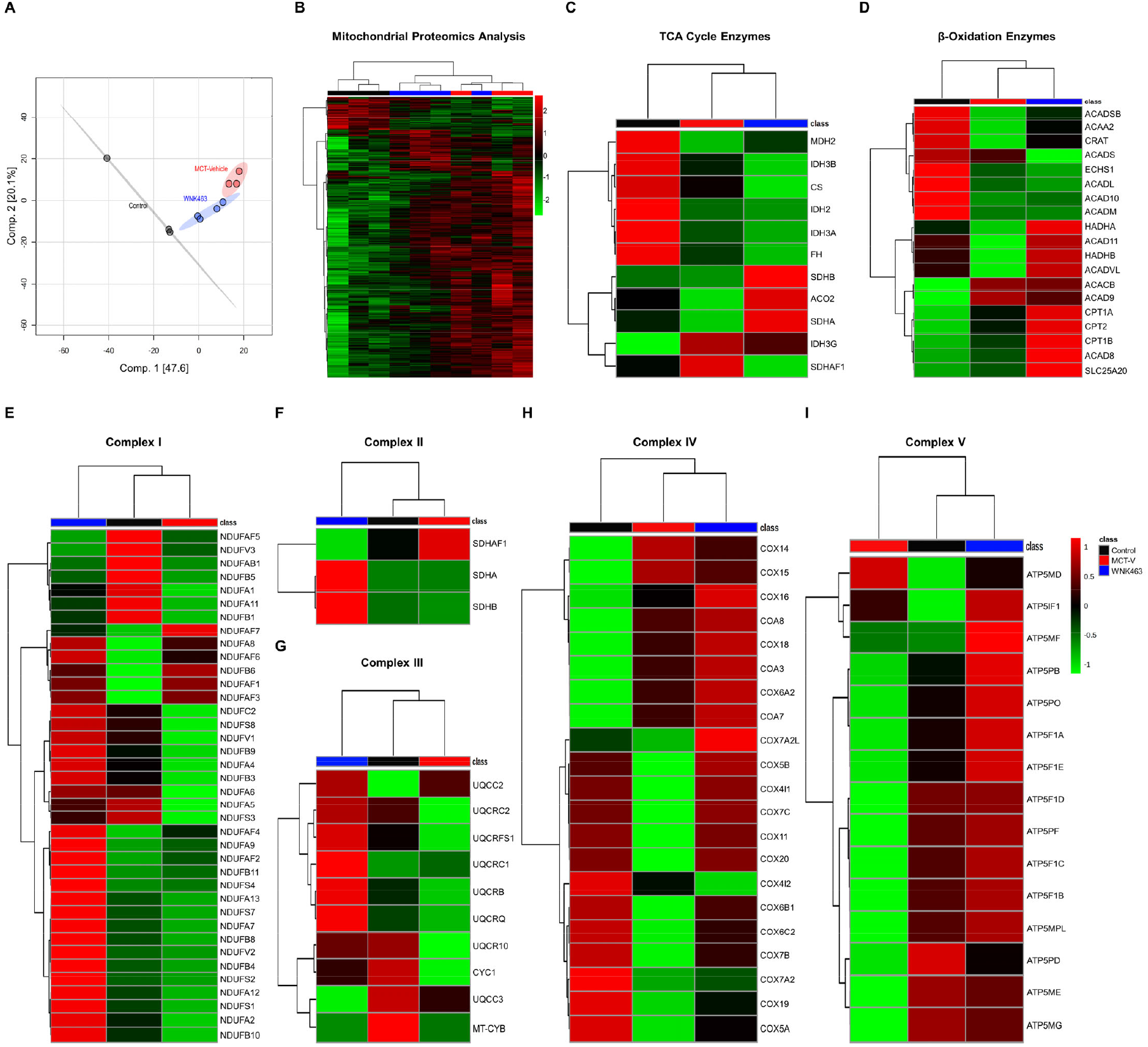
Quantitative Proteomics Reveals WNK463 Prevents Downregulation of Numerous Mitochondrial Metabolic Enzymes. (**A**) Principal component analysis and (**B**) hierarchical cluster analysis show WNK463 partially normalizes the global expression signature of mitochondrial/peroxisomal proteins. Hierarchical cluster analysis of (**C**) TCA cycle enzymes, (**D**) fatty acid β-oxidation enzymes, (**E**) complex I, (**F**) complex II, (**G**) complex III, (**H**) complex IV, and (**I**) complex V proteins demonstrate WNK463 partially corrects the dysregulation of multiple mitochondrial metabolic proteins.

### Metabolomics Profiling Demonstrated WNK463 Improved RV Metabolism

Global metabolomic profiling examined the metabolic state of the RV in control, MCT-V, and WNK463 rats. WNK463 modulated the RV metabolic signature, with a pattern that was an intermediate between control and MCT-V in hierarchical cluster analysis (**Figure 4A**). As discussed above, WNK463 prevented accumulation of pyruvate, suggesting less utilization of glycolysis (**Figure 1G**). We subsequently determined how WNK463 altered TCA cycle metabolites. As compared to controls, MCT-V RVs had elevated levels of nearly all TCA metabolites, which WNK463 combated (**Figure 4B**). Next, we analyzed 48 acylcarnitine-associated metabolites to probe mitochondrial FAO. Consistent with a previous study (22), nearly all acylcarnitines were reduced in MCT-V RVs. However, WNK463 increased concentrations of most acylcarnitines beyond that of control RVs (**Figure 4C**). In conclusion, these data revealed WNK463 altered RV metabolism. In particular, the restoration of metabolites in glycolysis, the TCA cycle, and acylcarnitines suggested multiple metabolic pathways were enhanced by WNK antagonism.

**Figure 4:**
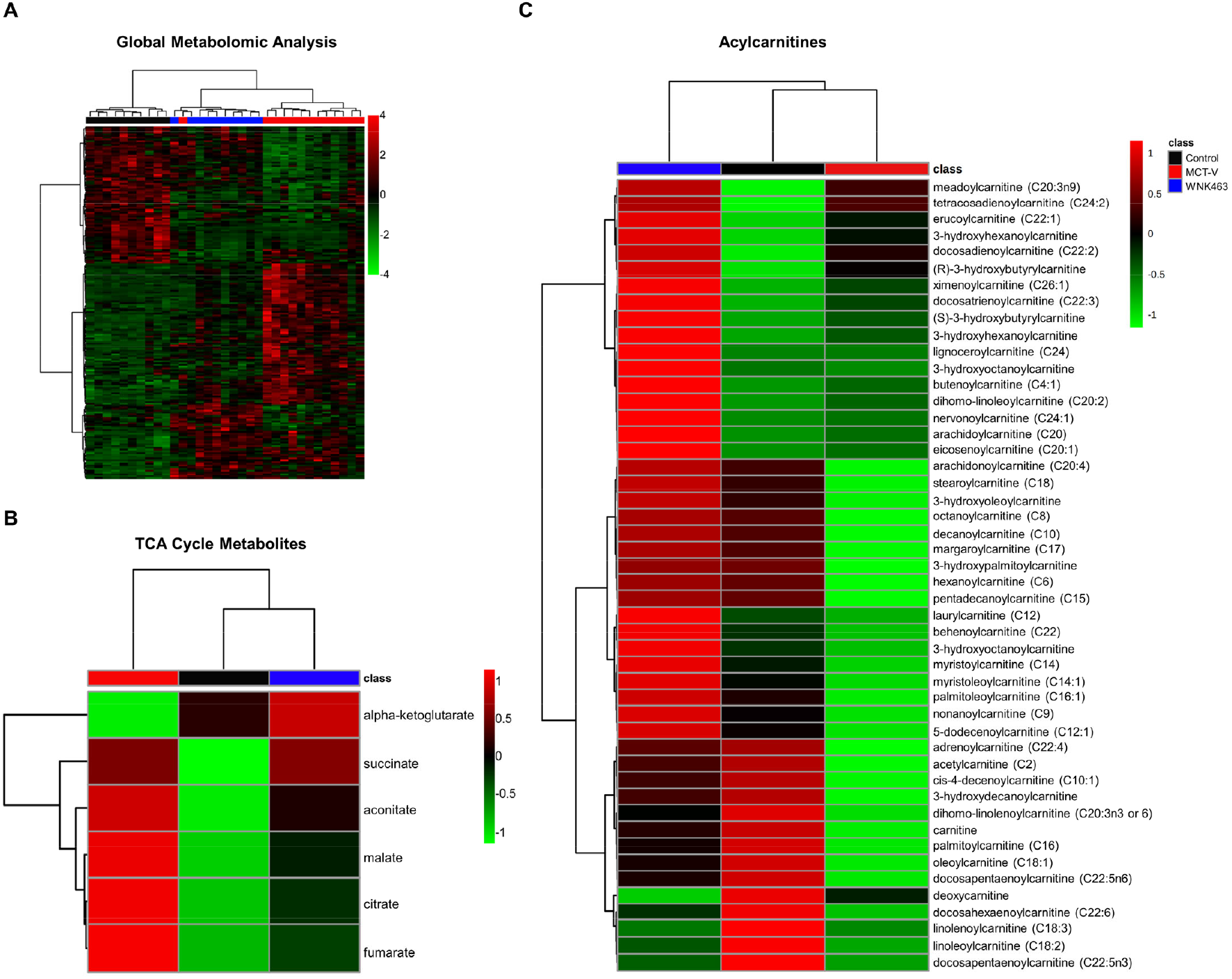
Global Metabolomics Profiling Shows WNK463 Partially Corrects RV Metabolic Derangements. Hierarchical cluster analysis of (**A**) global RV metabolomics data, (**B**) TCA cycle metabolites, and (**C**) acylcarnitines.

### WNK463 Mitigated Secondary Metabolic Effects of Impaired Mitochondrial Fatty Acid Oxidation

To determine what metabolic pathway was most disrupted in RVD, we integrated our proteomics and metabolomics data using joint pathway analysis. The three most dysregulated pathways included fatty acid degradation, the TCA cycle, and pyruvate metabolism (**Supplemental Figure 4**). Because other groups have also demonstrated disrupted lipid metabolism in RVD due to PAH (22-24), we focused our efforts on understanding how three secondary metabolic consequences of altered mitochondrial FAO (ω-fatty acid oxidation, glutaminolysis, and lipotoxicity/ceramide accumulation) (**Figure 5**) were impacted by WNK463. First, ω-fatty acid oxidation was increased in the MCT-V and MCT-WNK463 RVs as dicarboxylic fatty acid (DCA) levels were higher (**Figure 6A**) than control. However, MCT-WNK463 RVs had lower medium chain DCA levels when compared to MCT-V. In addition, our proteomics experiments revealed upregulation of many proteins responsible for ω-fatty acid oxidation and peroxisomal degradation of DCAs in both MCT-V and MCT-WNK463 RVs (**Figure 6B**). These data support the hypothesis that ω-fatty acid oxidation was induced by RV pressure overload, but the enhancement of mitochondrial FAO in MCT-WNK463 rats prevented accumulation of medium-chain DCAs. Second, we showed heightened glutaminolysis in the RVs of MCT-V rats as revealed by elevated levels of multiple glutaminolysis metabolites (glutamate, succinate, fumarate, malate, and pyruvate) (**Figure 6C**) and the mitochondrial glutaminolysis enzymes, glutaminase (GLS) and malic enzyme 2 (ME2) (**Figure 6D**). In contrast, WNK463 prevented the upregulation of GLS and ME2 and accumulation of several glutaminolysis metabolites (**Figure 6C-D**). Third, metabolomics profiling revealed increased abundance of eight species of ceramides, dihydroceramides, and hexosylceramides in the MCT-V RV, but WNK463 depressed concentrations of all eight lipids (**Figure 6E**). Finally, we observed ectopic lipid accumulation in MCT RVs, which WNK463 corrected (**Figure 6F**). In summary, these data suggest WNK463 combated accumulation of medium chain DCAs, glutaminolysis induction, and lipotoxicity.

**Figure 5:**
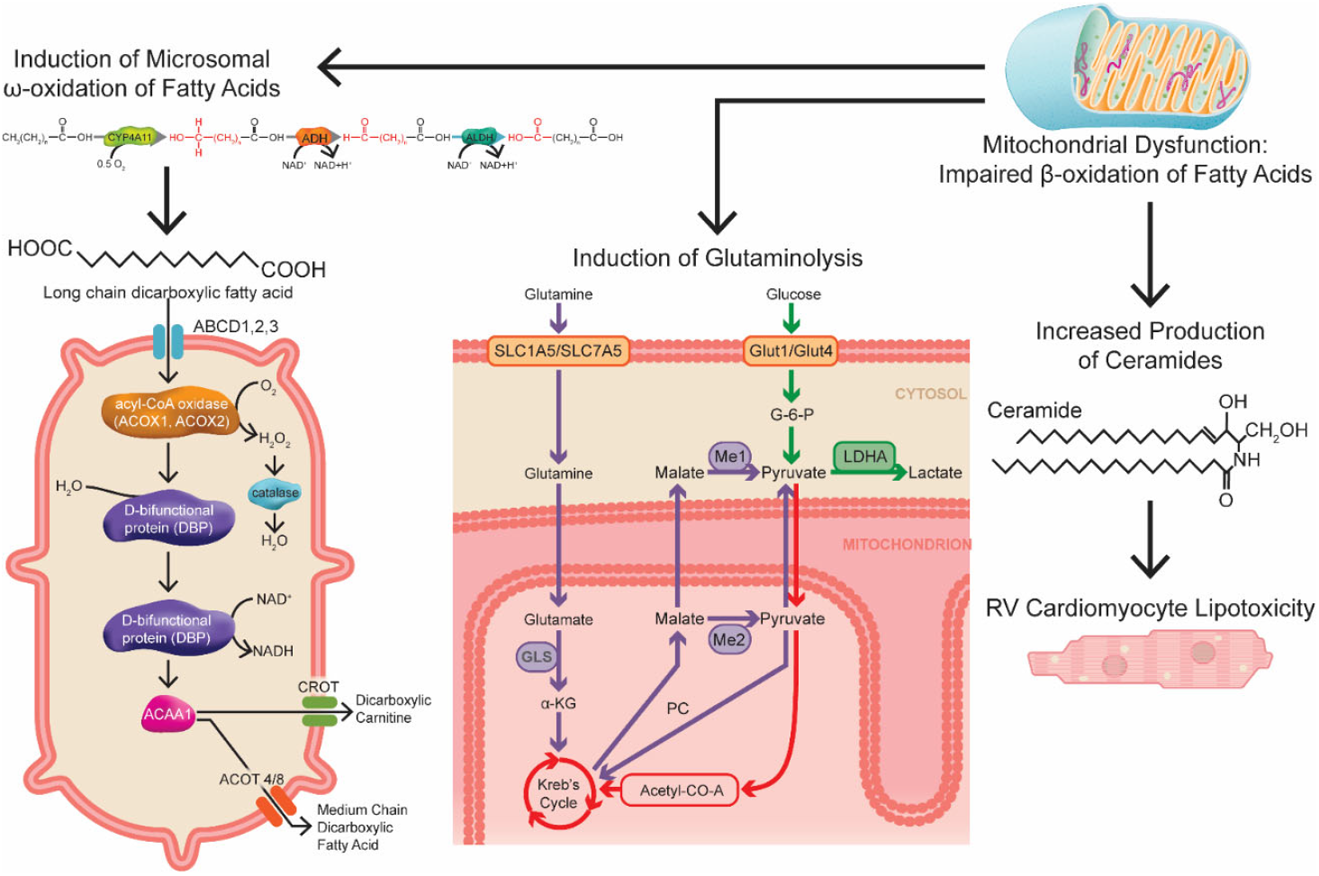
Diagram of the Metabolic Consequences of Impaired Mitochondrial β-Fatty Acid Oxidation. Three secondary metabolic consequences of altered mitochondrial FAO include induction of ω-fatty acid oxidation, glutaminolysis, and lipotoxicity due to ceramide accumulation.

**Figure 6:**
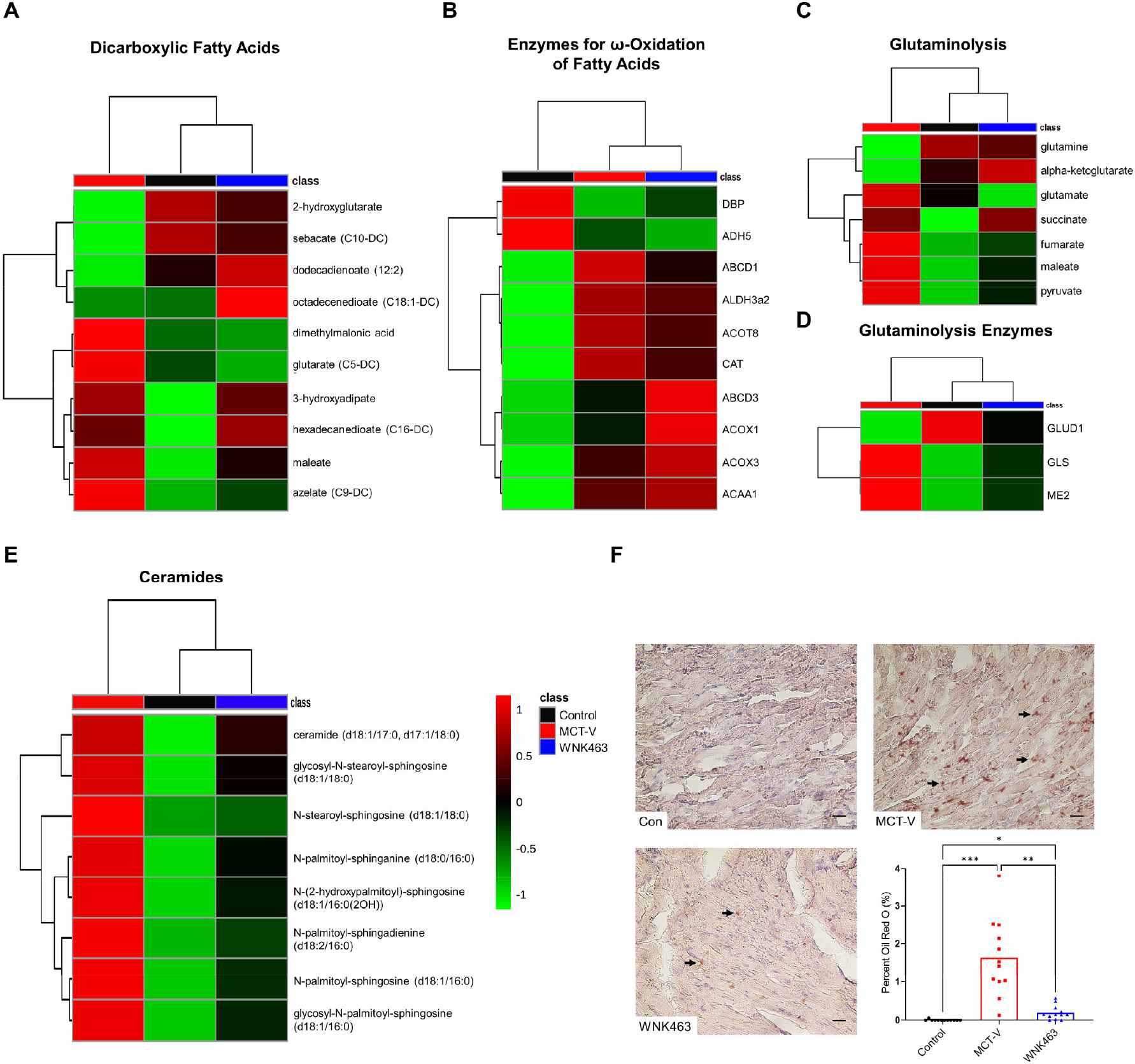
WNK463 Prevents Medium Chain Dicarboxylic Fatty Acid Accumulation, Reduces Glutaminolysis Induction, and Combats Lipotoxicity in the RV. WNK463 prevents accumulation of multiple dicarboxylic fatty acids (**A**), without modulating enzymes responsible for ω-oxidation of DCA (**B**). There are elevated levels of glutaminolysis intermediates (**C**) and glutaminolysis enzymes (**D**) in MCT-V RVs, which WNK463 normalizes. There is lipotoxicity in MCT-V as demonstrated by higher levels of multiple ceramide species (**E**) and RV intramyocardial lipid deposition as assessed by Oil Red O staining (**F**). Importantly, WNK463 counteracts these pathological changes (Control: 0.005±0.02%, MCT-V: 1.6±1.0%, WNK463: 0.2±0.2%, *p*=0.001 between MCT-V and WNK463, *n*=4 areas analyzed per RV and 3 RVs per group, 12 total RV areas analyzed per group). Scale bar 20 µm. \**p*<0.05, \*\**p*<0.01, and \*\*\**p*<0.001 by Brown-Forsythe and Welch ANOVA with Dunnett’s multiple comparisons test.

### WNK463 Did Not Alter Pulmonary Vascular Disease Severity

Next, we evaluated how WNK463 regulated pulmonary vascular disease severity to determine if our molecular changes were due to modulation of RV afterload. Echocardiography showed WNK463 did not significantly prolong pulmonary artery acceleration time (PAAT), an echocardiographic marker of pulmonary vascular disease severity (25), when compared to MCT-V (**Figure 7A**). PV loop analysis demonstrated WNK463 did not change RV afterload as RV systolic pressure (RVSP) (**Figure 7B**) and effective arterial elastance (Ea) (**Figure 7C**) were equivalent to MCT-V. Finally, pathological remodeling of the pulmonary arterioles was equivalent in MCT-V and MCT-WNK463 specimens (**Figure 7D-E**). Collectively, these data implied that the observed corrections in RV metabolism were not due to less severe pulmonary vascular disease.

**Figure 7:**
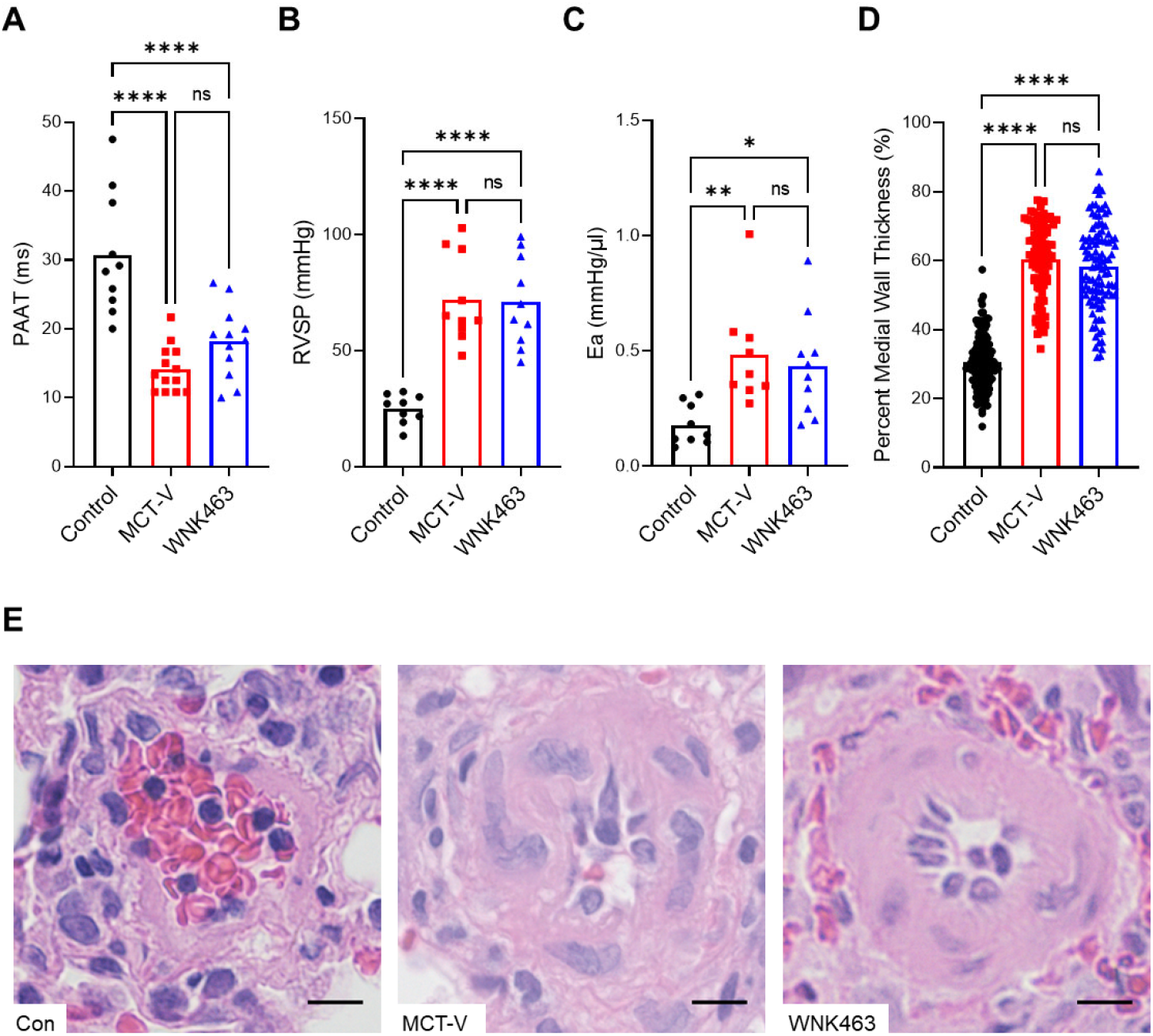
WNK463 Does Not Alter Pulmonary Vascular Disease Severity. (**A**) There is no statistical difference in pulmonary artery acceleration time (PAAT) between MCT-V and WNK463 as assessed by echocardiography (Control: 31±9, MCT-V: 14±3, WNK463: 18±5 ms, *p*=0.22 between MCT-V and WNK463, *n*=10 control, 13 MCT-V, and 12 WNK463). (**B**) No change in RV systolic pressure (RVSP) between MCT-V and WNK463 (Control: 25±6, MCT-V: 72±19, WNK463: 71±19 mmHg, *p*=0.99 between MCT-V and WNK463, *n*=9 control, 10 MCT-V, and 10 WNK463). (**C**) No difference in effective arterial elastance (Ea) between MCT-V and WNK463 (Control: 0.18±0.09, MCT-V: 0.48±0.22, WNK463: 0.43±0.22 mmHg/µl, *p*=0.84 between MCT-V and WNK463, *n*=9 control, 9 MCT-V, and 10 WNK463). (**D**) Histologically, there was no change in pulmonary small arteriole remodeling between MCT-V and WNK463 (Control: 31±8%, MCT-V: 60±11%, WNK463: 58±12% medial wall thickness, *p*=0.52, *n*=134 control, 94 MCT-V, and 105 WNK463 pulmonary arterioles from 4 lung tissues per group). **(E)** Representative H&E images of pulmonary arterioles. Scale bar 10 µm. \**p*<0.05, \*\**p*<0.01, \*\*\*\**p*<0.0001, and (ns) not significant by one-way ANOVA with Tukey’s multiple comparisons test in (**A-C**) or Brown-Forsythe and Welch ANOVA with Dunnett’s multiple comparisons test in (**D**).

### WNK463 Decreased RV Hypertrophy and Fibrosis, and Enhanced RV Systolic and Diastolic Function

WNK463 decreased the Fulton index (ratio of RV to LV and septum weight) (**Figure 8A**) and ratio of RV free wall weight to total body weight (**Figure 8B**). Furthermore, WNK463 reduced RV cardiomyocyte cross sectional area, but it was not completely normalized (**Figure 8C-D**). Thus, WNK463 mitigated pathological RV hypertrophy.

**Figure 8:**
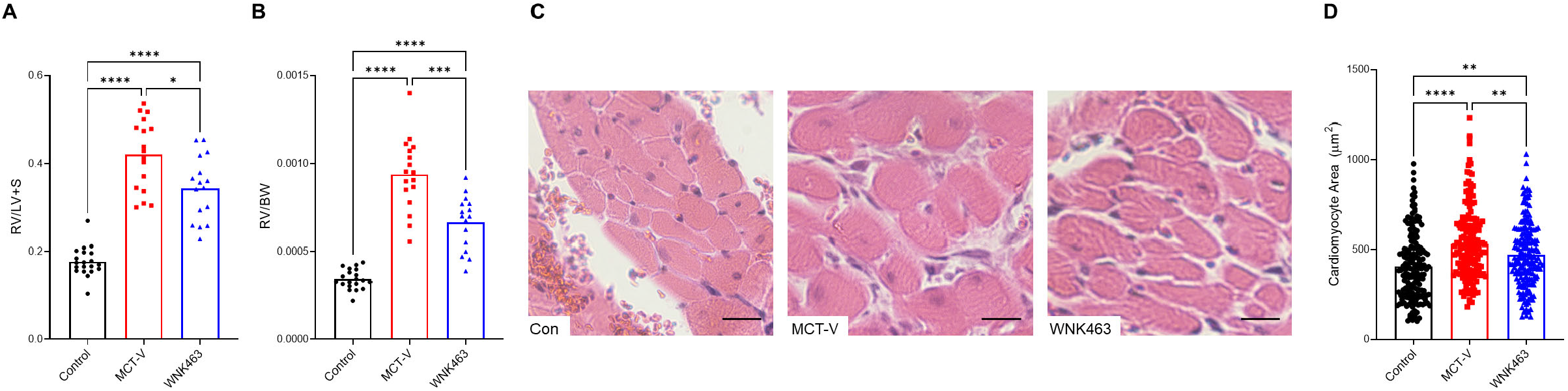
WNK463 Combats Pathological Right Ventricular Hypertrophy. Compared to MCT-V, WNK463 reduces (**A**) the Fulton index (Control: 0.18±0.03, MCT-V: 0.42±0.08, WNK463: 0.34±0.07, *p*=0.02 between MCT-V and WNK463, *n*=20 control, 17 MCT-V, and 16 WNK463), (**B**) RV weight normalized to body weight (Control: 0.0003±0.00006, MCT-V: 0.0009±0.0002, WNK463: 0.0007±0.0002, *p*=0.0004 between MCT-V and WNK463, *n*=20 control, 17 MCT-V, and 16 WNK463), and cardiomyocyte cross-sectional area (Control: 408±186, MCT-V: 530±200, WNK463: 470±180 µm^2^, *p*=0.007 between MCT-V and WNK463, *n*=184 control, 177 MCT-V, and 196 WNK463 cardiomyocytes measured from 3 RVs per group) with representative images in (**C**) and quantification in (**D**). Scale bar 20 µm. \**p*<0.05, \*\**p*<0.01, \*\*\**p*<0.001 and \*\*\*\**p*<0.0001 by Brown-Forsythe and Welch ANOVA with Dunnett’s multiple comparisons test in (**A-B**) or one-way ANOVA with Tukey’s multiple comparisons test in (**D**).

We then used echocardiography and PV loops to examine RV function. Echocardiographic analysis demonstrated WNK463 increased tricuspid annular plane systolic excursion (TAPSE), percent RV free wall thickening, cardiac output, and cardiac output normalized to body mass (**Figure 9A-D**) as compared to MCT-V. PV loop analysis showed end-systolic elastance/effective arterial elastance (Ees/Ea), the gold standard of RV function (26), was higher in MCT-WNK463 than MCT-V (**Figure 9E**). In addition to the augmented RV systolic function, WNK463 treatment enhanced RV diastolic function as determined by a reduction in RV end-diastolic pressure (RVEDP) and RV τ (relaxation time) (**Figure 9F-G**).

**Figure 9:**
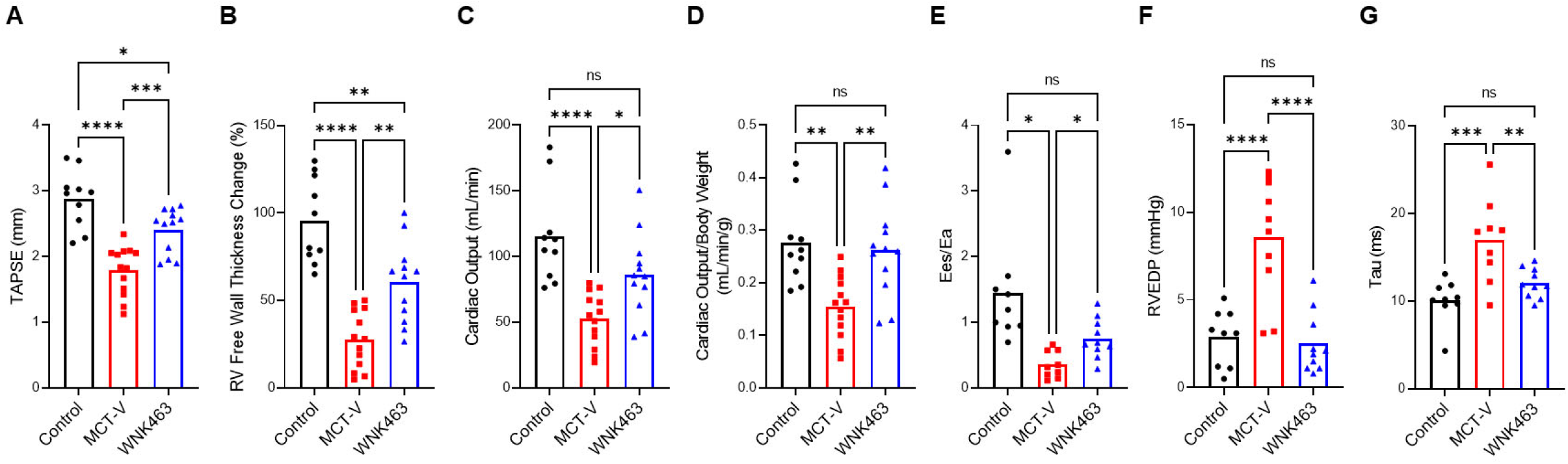
WNK463 Enhances RV Systolic and Diastolic Function. WNK463 increases (**A**) TAPSE (Control: 2.9±0.4, MCT-V: 1.8±0.4, WNK463: 2.4±0.3 mm, *p*=0.0009 between MCT-V and WNK463, *n*=10 control, 13 MCT-V, and 12 WNK463), (**B**) percent RV free wall thickness change (Control: 96±24%, MCT-V: 28±17%, WNK463: 60±23%, *p*=0.001 between MCT-V and WNK463, *n*=10 control, 13 MCT-V, and 12 WNK463), (**C**) cardiac output (Control: 115±36, MCT-V: 52±20, WNK463: 86±31 mL/min, *p*=0.02 between MCT-V and WNK463, *n*=10 control, 13 MCT-V, and 12 WNK463), and (**D**) cardiac output normalized to body weight (Control: 0.28±0.08, MCT-V: 0.15±0.06, WNK463: 0.26±0.09 mL/min/g, *p*=0.004 between MCT-V and WNK463, *n*=10 control, 13 MCT-V, and 12 WNK463) as assessed by echocardiography. WNK463 augments RV-PA coupling (Ees/Ea) (Control: 1.4±0.9, MCT-V: 0.4±0.2, WNK463: 0.8±0.3, *p*=0.01 between MCT-V and WNK463, *n*=9 control, 9 MCT-V, and 10 WNK463) (**E**), reduces RVEDP (Control: 3±2, MCT-V: 9±3, WNK463: 3±2 mmHg, *p*<0.0001 between MCT-V and WNK463, *n*=9 control, 10 MCT-V, and 10 WNK463) (**F**), and improves RV diastolic function (tau) (Control: 10±2, MCT-V: 17±5, WNK463: 12±2 ms, *p*=0.008 between MCT-V and WNK463, *n*=9 control, 9 MCT-V, and 10 WNK463) (**G**) when evaluated by closed-chest PV loop analysis. \**p*<0.05, \*\**p*<0.01, \*\*\**p*<0.001, \*\*\*\**p*<0.0001, and (ns) not significant by one-way ANOVA with Tukey’s multiple comparisons test in (**A-D**) and (**F-G**) or Brown-Forsythe and Welch ANOVA with Dunnett’s multiple comparisons test in (**E**).

### Hypochloremia was Associated with Exacerbated RV Dysfunction in PAH

Finally, we analyzed the effects of hypochloremia, a condition at activates WNK1 (17), on RV function in a cohort of 217 PAH patients (**Supplemental Table 2**). First, we plotted the relationship between right atrial (RA) pressure and pulmonary vascular resistance (PVR), and found hypochloremic patients had higher RA pressure at all PVR values than normochloremic patients (**Figure 10A**). Furthermore, as PVR increased, hypochloremic patients had a more rapid decline in cardiac output (**Figure 10B**). These data are consistent with the hypothesis that hypochloremia is associated with more severe RV dysfunction in PAH patients.

**Figure 10:**
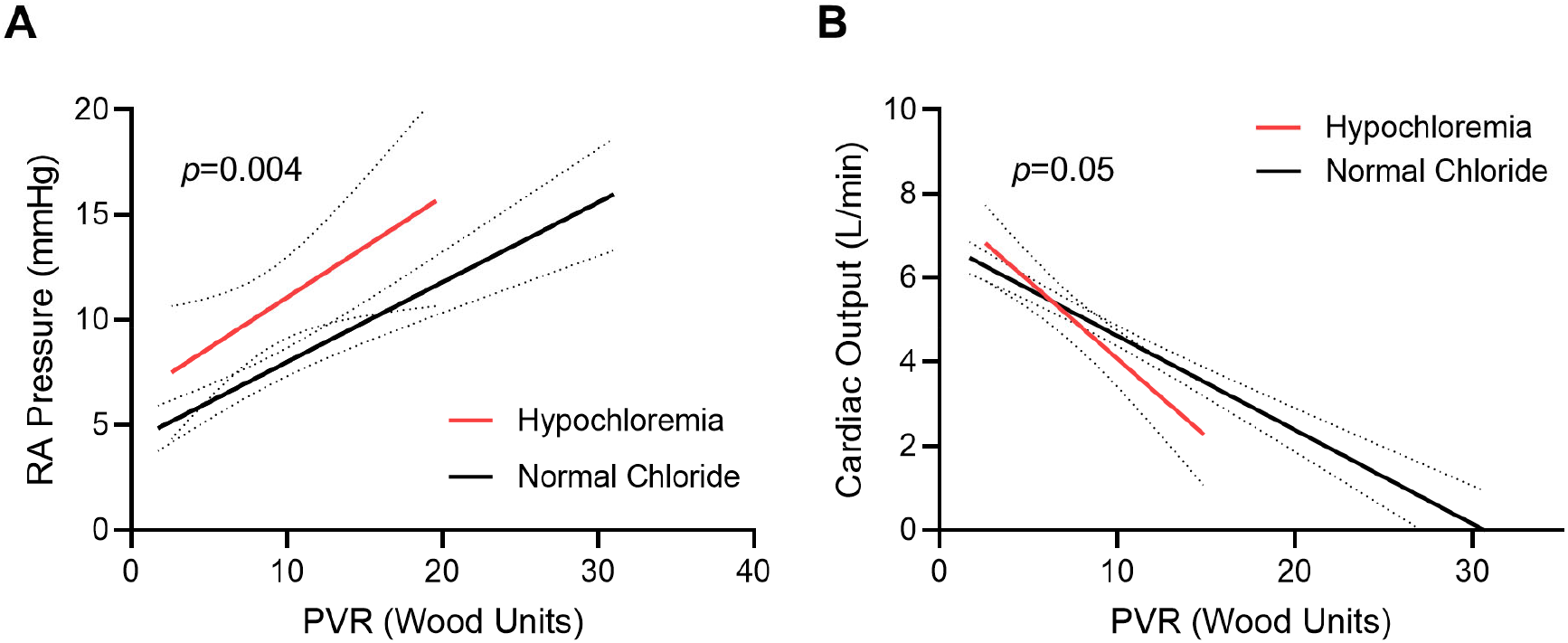
Hypochloremia Results in Exaggerated RV Dysfunction in PAH. (**A**) PAH patients with hypochloremia (serum chloride ≤101 mmol/L) have higher RA pressures at every pulmonary vascular resistance (PVR) as compared to PAH patients with normal serum chloride levels (*p*=0.0004 between y-intercepts, *p*=0.58 between slopes, *n*=38 patients with hypochloremia and 162 with normal chloride). (**B**) PAH patients with hypochloremia have a more rapid descent in cardiac output as PVR increases (*p*=0.05 between slopes, *p*=0.56 between y-intercepts, *n*=24 patients with hypochloremia and 131 patients with normal chloride).

## Discussion

In this study, we show small molecule inhibition of WNK1 signaling prevents upregulation of GLUT1 and GLUT4 via mitigation of AS160 phosphorylation, which subsequently restores the levels of glucose metabolites in the RV. The normalization of RV glucose uptake by WNK463 depresses excess protein O-GlcNAcylation and glycation, and restores levels of mitochondrial enzymes involved in the TCA cycle, FAO, and ETC complexes. These proteomic changes are matched with metabolic shifts on metabolomics analysis that are indicative of partial correction of RV metabolism. Integration of our proteomics and metabolomics analyses identifies FAO as the most altered pathway in preclinical RVD. WNK463 combats the secondary consequences of impaired mitochondrial FAO in the dysfunctional RV as there is less glutaminolysis induction and accumulation of medium chain DCAs and ceramides. Importantly, these molecular changes result in enhanced RV systolic and diastolic function, which is not due to alteration in pulmonary vascular disease severity. Finally, hypochloremia in PAH results in more severe RVD, providing evidence that this pathway may be relevant in human disease. In summary, these data identify WNK1 as a druggable target for PAH-associated RVD due to modulation of glucotoxicity and subsequent metabolic derangements.

WNK463 prevents both protein O-GlcNAcylation and glycation, which results in less mitochondrial protein dysregulation and augmented metabolic function. Our results are congruent with several other studies showing O-GlcNAcylation and glycation directly promote mitochondrial dysfunction. Importantly, there is evidence that these PTMs modulate mitochondrial function through multiple mechanisms. First, several mitochondrial proteins are O-GlcNAcylated (27-29) and glycated (30), and these PTMs can directly alter enzymatic activity. In addition, cardiac-specific transgenic overexpression of O-linked β-*N*-acetylglucosamine transferase (OGT), the enzyme that catalyzes the addition of O-GlcNAc, decreases mRNA levels of enzymes in FAO, the TCA cycle, and oxidative phosphorylation, which ultimately depresses mitochondrial function (31). Thus, excess O-GlcNAcylation may directly modulate enzymatic activity via PTM and depress transcription of enzymes in multiple mitochondrial metabolic pathways, which in total causes metabolic dysregulation. With regards to protein glycation, knockout of DJ-1, the protein that reverses protein glycation (32), reduces cardiomyocyte mitochondrial DNA content resulting in cardiomyopathy (13), suggesting protein glycation may alter mitochondrial biogenesis and/or stability. Moreover, overexpression of DJ-1 mitigates excess protein glycation which prevents the development of ischemia-reperfusion-induced LV failure (12). Thus, our results and other existing data show O-GlcNAcylation and glycation adversely affect mitochondrial metabolic function and cardiac function.

Our data may provide a molecular explanation of the increased mortality associated with hypochloremia in PAH (14,15) and in left heart failure patients (33-35). Based on our findings, we propose hypochloremia activates WNK1, which subsequently heightens glucose uptake resulting in excess protein O-GlcNAcylation and glycation. These pathological PTMs cause mitochondrial metabolic dysfunction, which then increases the demand for glycolytic metabolism and cardiomyocyte glucose uptake, ultimately resulting in a vicious downward cycle culminating in cardiac failure. However, WNK1-mediated glucotoxicity may be particularly important in RV failure because the Human Cardiac Cell Atlas reveals RV cardiomyocytes have higher expression levels of WNK1, GLUT4, and AS160 than LV cardiomyocytes (specifically, in cardiomyocyte population 2, which is more enriched in the RV than LV) (36) (Supplemental Table 2). Additionally, RV cardiomyocytes exhibit higher rates of glycolysis than LV cardiomyocytes (37), which suggests alterations in glucose metabolism may have heightened consequences in the RV.

Our findings also provide important insights into the interplay of multiple metabolic pathways implicated in RV dysfunction in PAH. First, we show WNK463 restores FAO in the RV marked by increased levels of multiple acylcarnitine species (**Figure 4C**). Interestingly, RV acylcarnitines are reduced in human PAH (22), which suggests our results are directly relevant to human disease. Furthermore, we demonstrate ceramide accumulation in RV failure, a finding also observed in human PAH (22,23). Moreover, our data show ω-fatty acid oxidation is accentuated in rodent RVD (**Figure 6B**). Human studies also suggest ω-fatty acid oxidation is induced in PAH as octadecanedioate, a long chain DCA, is elevated in PAH patients (38). Moreover, levels of octadecanedioate are inversely associated with RV function in pediatric PAH (39), providing further support that RVD is associated with ω-fatty acid oxidation. Finally, we show glutaminolysis is induced in RVD, but corrected with WNK463. Interestingly, inhibition of glutaminolysis enhances RV function in preclinical PAH, and there are higher levels of SLC1A5, the protein that imports glutamine into the cell, in human PAH RV specimens (40). In summary, we and others identify multiple metabolic derangements in the failing RV, and there appears to be a highly-interdependent relationship between defective mitochondrial FAO, induction of ω-fatty acid oxidation and glutaminolysis, and lipotoxicity. Importantly, disturbances in all of these metabolic pathways are observed in human studies, suggesting our results have direct human relevance.

Peroxisomes are secondary metabolic organelles that are responsible for metabolism of long chain and complex fatty acids (41), but the role of peroxisomes in cardiac dysfunction is understudied. The importance of peroxisomes in proper cardiac function is demonstrated by cardiomyopathy caused by cardiomyocyte-specific knockout of peroxisome proliferator-activated receptor gamma (PPARϒ) (42). Moreover, the PPARϒ agonist, pioglitazone enhances FAO in isolated RV cardiomyocytes and rescues RVD in Sugen-hypoxia rats (6), albeit in the setting of reduced PAH severity. Our proteomics analysis reveals an increase in multiple peroxisomal proteins in RV failure (**Supplemental Figure 5**), suggesting compensatory peroxisome biogenesis occurs in RV pressure overload. Surprisingly, multiple peroxisomal proteins are lower in abundance in the RV than the LV (41). This implies there may be chamber-specific differences in peroxisomal importance, and that the RV may have less of a peroxisomal reserve than the LV. However, future studies are needed to clearly delineate the role of peroxisomes in RVD.

Although the rationale to inhibit WNK1 signaling in our study was based on a clinical observation, other molecular mechanisms likely promote WNK1 upregulation in RVD. First, WNK1 levels are increased by tumor necrosis factor-α (TNFα) via reducing expression of NEDD4-2 E3-ligase (neuronal precursor cell-expressed developmentally downregulated 4-2 E3-ubiquitin ligase), a protein that degrades WNK1 in kidney cells (43). This is directly relevant to RVD because TNFα levels increase with the severity of RVD in rodent PAH (44). Moreover, aldosterone post-transcriptionally enhances WNK1 expression via miR-192 in the kidney (45). Again, this may be pertinent to RVD from PAH as serum aldosterone levels are inversely associated with cardiac output in human PAH (46). Thus, multiple pathways likely converge to promote WNK1 upregulation/activation and subsequent glucotoxicity in RVD, which may explain why WNK inhibition was so efficacious.

### Study Limitations

Our study has important limitations that must be acknowledged. First, all of our animal studies were performed in male MCT rats because we wanted to use the most severe model of RV failure to probe WNK1 signaling. Moreover, we only used one model of PAH and RV failure, but as described above, many of the metabolic disturbances we documented are also present in human analyses, so we believe this model is directly relevant to human RV failure. The beneficial effects of WNK463 could be due to inhibition of other isoforms of WNK, but we believe it is predominately mediated by WNK1 as we were unable to detect WNK2 in cardiac extracts (**Supplemental Figure 6**). This is consistent with low WNK2 cardiac abundance observed in the Human Protein Atlas (47). Additionally, WNK3 and WNK4 mRNA is not detected in cardiac tissue (17), further supporting the hypothesis that WNK1 is the most important WNK isoform for cardiac physiology. There may have been differences in mitochondrial robustness leading to variability in efficacy extraction in our proteomics analysis. However, that is less likely as most of the mitochondrial proteins were actually higher in the diseased animals, including multiple mitochondrial membrane and ribosomal proteins (**Supplemental Figure 7**). The reduction in RVEDP with WNK463 could be due to the diuretic effects of the compound (48), but the normalization of RV tau suggests a true change in RV diastolic function. Finally, the increased abundance of HBP intermediates may impact RV physiology independent of protein O-GlcNAcylation as UDP-GlcNAc is used to synthesize extracellular matrix components (49). Consistent with this possibility, we detected a reduction in RV fibrosis in WNK463 treated rats (**Supplemental Figure 8**), which may be another mechanism underlying improvements in RV systolic and diastolic function with WNK463.

### Perspectives

#### Competency in Medical Knowledge

Low chloride is associated with increased mortality in PAH, but a mechanistic explanation is lacking. Here, we show inhibition of the chloride-activated protein, WNK1 restores RV function by modulating glucotoxicity and fatty acid metabolism in rodent PAH.

#### Translational Outlook

Our data identify WNK1 as a potential druggable target for PAH-associated RVD. Further studies examining the safety and tolerability of WNK inhibition are needed to determine if our results could be translated to PAH patients with RV failure.

## Abbreviations List

Aco2: Aconitase 2
AS160: 160 kDa substrate of the Akt serine/threonine kinase
ATP: Adenosine triphosphate
DCA: Dicarboxylic fatty acid
DJ-1: Protein deglycase
Ea: Effective arterial elastance
Ees: End-systolic elastance
ETC: Electron transport chain
FAO: Fatty acid oxidation
GFAT1: Glutamine-fructose-6-phosphate transaminase 1
GLO1: Glyoxalase 1
GLO2: Glyoxalase 2
GLS: Glutaminase
GLUT1: Glucose transporter 1
GLUT4: Glucose transporter 4
LV: Left ventricle/ventricular
MCT: Monocrotaline
MCT V: Monocrotaline-vehicle
ME2: Malic enzyme 2
OGA: O-GlcNAcase
OGT: O-linked β-*N*-acetylglucosamine transferase
PAAT: Pulmonary artery acceleration time
PAH: Pulmonary arterial hypertension
PPARγ: Peroxisome proliferator-activated receptor gamma
PTM: Post-translationally modify/modifications
PV: Pressure-volume
PVR: Pulmonary vascular resistance
RA: Right atrial
RV: Right ventricle/ventricular
RVEDP: Right ventricular end-diastolic pressure
RV-PA: Right ventricular-pulmonary artery
RVD: Right ventricular dysfunction
RVSP: Right ventricular systolic pressure
Sdha: Succinate dehydrogenase complex flavoprotein subunit A
Sdhb: Succinate dehydrogenase complex iron sulfur subunit B
TAPSE: Tricuspid annular plane systolic excursion
Tau/τ: Right ventricular relaxation time
TCA: Tricarboxylic acid
TNFα: Tumor necrosis factor-α
UDP-GlcNAC: Uridine diphosphate *N*-acetylglucosamine
WNK: <u>W</u>ith <u>N</u>o <u>L</u>ysine kinase

## Author Contributions

SZP and KWP designed research studies, conducted experiments, acquired data, analyzed data, and prepared the manuscript. ME conducted experiments, acquired data, and analyzed data. TT acquired and analyzed data. All authors revised the manuscript and approved the submitted version.

## Acknowledgments

Echocardiography and confocal and electron microscopy imaging were completed at the University Imaging Center. We thank the University of Minnesota Histology and Research Laboratory in the Clinical and Translational Science Institute for their assistance with processing lung histology. We thank Drs. LeeAnn Higgins and Todd Markowski of the University of Minnesota Center for Mass Spectrometry and Proteomics for their assistance in obtaining the quantitative mass spectrometry data. We also thank Cynthia Faraday for her assistance with figure design.

