## Supplemental Data for "With No Lysine Kinase 1 Promotes Right Ventricular Dysfunction Via Glucotoxicity"

**Supplemental Methods**

*Animal Models and Treatment*. Male Sprague Dawley rats (200-250 g; 7-8 weeks old) (Charles River Laboratories, Wilmington, MA) received a single subcutaneous injection of monocrotaline (MCT) (60 mg/kg) (Sigma-Aldrich, St. Louis, MO) or phosphate-buffered saline (control). Two weeks after MCT injection, rats were given daily intraperitoneal injections of either 3 mg/kg WNK463 (D&C Chemicals, Wanze, Belgium) or vehicle (5% propylene glycol, 0.475% Pluronic, 0.475% Klucel LF, and 94.05% water) (48) for 10 days. We estimated a sample size of 15-20 rats per group to reach statistical significance. Rats were randomized to treatment groups. All animal studies were approved by the University of Minnesota Institutional Animal Care and Use Committee.

*Western Blot Analysis.* Immunoblots with 25 µg of RV protein extracts were imaged with the Odyssey Infrared Imaging system (Lincoln, NE) as previously described (20). Post transfer SDS-PAGE gels were stained with Coomassie brilliant blue (CBB) and the band corresponding to the myosin heavy chain was used as the loading control (20,50). Antibodies used in this study are listed in **Supplemental Table 1**. Complete Western blot images are shown in **Supplemental Figure 1**.

*Immunofluorescence and Confocal Microscopy*. RV free wall tissues were cryosectioned at 10-µm. Antigen retrieval buffer (10 µM Tris, 1 mM EDTA, and 0.05% Tween 20) was applied to all sections while the slides were heated to 100ºC for 5 minutes. Sections were then blocked in 5% goat serum (Thermo Fisher, Waltham, MA) twice for 5 minutes and incubated in primary antibody for 72 hours. Sections were subsequently incubated in fluorescent-dye conjugated secondary antibody for 1-2 hours and then in TrueVIEW Autofluorescence Quenching Kit with DAPI (Vector Laboratories, Burlingame, CA) per manufacturer's protocol. Slides were mounted in Anti-Fade Reagent (Molecular Probes, Eugene, OR). Antibodies used for immunofluorescence are listed in **Supplemental Table 1**. Images were obtained with an Olympus FV1000 BX2 upright confocal microscope (Tokyo, Japan) at the University of Minnesota Imaging Center.

*Mitochondrial/Peroxisome Enrichments.* RV mitochondrial/peroxisome enrichments were isolated with a mitochondrial isolation kit (Abcam, Cambridge, MA) as previously described (51). Then, the mitochondrial/peroxisome pellet fraction was resuspended in lysis buffer and supplemented with protease inhibitor (Thermo Fisher). The samples were briefly sonicated with a Branson Digital Sonifier 250 (Branson Ultrasonics, Brookfield, CT) and then underwent pressure cycling (alternating 35 KPSI for 20 seconds and 0 KPSI for 10 seconds for 60 cycles at 37ºC). Protein concentration was determined by Bradford assay.

*Quantitative Mass Spectrometry.* A 25 µg aliquot of each sample obtained in the mitochondrial/peroxisomal enrichment was diluted in half with lysis buffer and then this sample was then diluted 5-fold with water. Trypsin (Promega, Madison, WI) was added in a 1:40 ratio. Samples were incubated overnight at 37ºC and after incubation, frozen at -80ºC and vacuum dried. Each sample was cleaned with a 1 cc Waters Oasis MCX cartridge (Waters Corporation, Milford, MA), eluates were vacuum dried and resuspended in 100 µl 0.1 M triethylammonium bicarbonate, pH 8.5 to a final concentration of 1 µg/µl. For each channel in the TMT10plex™, a 20 µg aliquot of the sample was made and labeled with 0.2 mg TMT10plex™ Isobaric Label Reagent (Thermo Scientific, Waltham, MA) per the manufacturer's protocol. After TMT labeling, all samples were multiplexed together.

The TMT™ sample was then dried *in vacuo*, cleaned with a 1 cc Waters Oasis MCX solid phase extraction cartridge (Waters Corporation), and dried *in vacuo* again. The sample was resuspended in 20 mM ammonium formate, pH 10 in 98:2 water:acetonitrile (Buffer A) and fractioned by high pH C18 reversed-phase (RP) chromatography (52) with the following changes. A Shimadzu Prominance HPLC (Shimadzu, Columbia, MD) with a Hot Sleeve 2 L Column Heater (Analytical Sales and Services, Inc., Flanders, NJ) was used with a SecurityGuard precolumn housing a Gemini NX-C18 cartridge (Phenomenex, Torrance, CA) attached to a C18 XBridge column (150 mm x 2.1 mm internal diameter, 5 µm particle size) (Waters Corporation). Buffer A was 20 mM ammonium formate, pH 10 in 98:2 water:acetonitrile and Buffer B was 20 mM ammonium formate, pH 10 in 10:90 water:acetonitrile. Flow rate was 200 µl/min with a gradient from 2-7% Buffer B over 0.5 minutes, 7-15% Buffer B over 7.5 minutes, 15-35% Buffer B over 45 minutes, and 35-60% Buffer B over 15 minutes. Fractions were collected every 2 minutes and UV absorbances were monitored at 215 and 280 nm.

*Global Metabolomics Profiling.* RV metabolomics profiling on frozen RV free wall specimens was completed by Metabolon, Inc. (Durham, NC). In brief, ultrahigh performance liquid chromatography/electrospray ionization tandem mass spectrometry was performed on a Waters ACQUITY ultra-performance liquid chromatography (UPLC) and a Thermo Scientific Q-Exactive high resolution/accurate mass spectrometer interfaced with a heated electrospray ionization (HESI-II) source and Orbitrap mass analyzer. Semi-quantitative metabolite values are reported.

*Lung Histology.* Lung tissues were fixed in 10% formalin, embedded in paraffin, sectioned at 4-μm, and stained with H&E by the University of Minnesota Histology and Research Laboratory in the Clinical and Translational Science Institute. Images were acquired on a Zeiss AxioCam IC (Oberkochen, Germany) and measurements were completed in FIJI. Percent medial thickness of small pulmonary arterioles was calculated as 100 times (outer diameter-inner diameter)/outer diameter (20).

*Cardiac Histology.* RV tissues were fixed in 10% formalin, embedded in paraffin, sectioned at 10-μm, and stained with H&E by the University of Minnesota Histology and Research Laboratory in the Clinical and Translational Science Institute to evaluate cardiomyocyte area. RV cryosections at 10-μm were stained with trichrome stain (Abcam) to assess fibrosis. Percent fibrosis was assessed by measuring the area of tissue stained blue divided by total tissue area. All measurements were completed in FIJI (Bethesda, MD).

*Oil Red O Stain*. RV cryosections at 10-μm were stained with 0.5% Oil Red O (Sigma-Aldrich) in propylene glycol for 4 hours and counterstained with hematoxylin (Abcam, Cambridge, MA) to identify intramyocardial lipid deposits. Sections were imaged with a Zeiss AxioCam IC and percent Oil Red O stain was assessed with FIJI.

*Rodent Echocardiography*. Echocardiography was completed with a 37.5-MHz transducer (VisualSonics) and Vevo2100 ultrasound system as described previously (20). M-mode and 2-D images were obtained to measure tricuspid annular plane systolic excursion (TAPSE) and RV free wall thickness during end diastole and end systole. Pulsed-wave Doppler assessed pulmonary artery flow velocity time integral (25). Representative images obtained during echocardiography are in **Supplemental Figure 2**.

*Closed-Chest Pressure Volume Loop Analysis.* Rats were induced with 5% isoflurane and then maintained on 2-3% isoflurane. They were ventilated with the SomnoSuite small animal anesthesia system (Kent Scientific, Torrington, CT). A high-fidelity catheter (Scisense 1.9F pressure-volume, Transonic Systems, Ithaca, NY) was placed into the RV via the right internal jugular vein and right atria. RV pressure and volume were measured with the Transonic AV500 Pressure-Volume Measurement System and then analyzed on LabScribe version 4 (iWorx Systems, Dover, NH). Representative PV loops images are shown in **Supplemental Figure 3**.

*PAH Patient Cohort.* PAH patients from the University of Minnesota Pulmonary Hypertension Program (53) were assessed. PAH was defined as mean pulmonary arterial pressure (mPAP) ≥20 mmHg, pulmonary capillary wedge pressure ≤15 mmHg, and pulmonary vascular resistance (PVR) ≥3.0 Wood units not due to other causes such as left-sided heart disease, chronic lung disease, or chronic thromboembolic disease as determined by echocardiography, pulmonary function tests, computed tomography imaging, ventilation perfusion scans, or invasive pulmonary angiogram (54). Serum chloride level closest to the right heart catheterization data was evaluated (average time difference 23 ± 40 days). Hypochloremia was defined as serum chloride ≤101 mmol/L as we previously defined (15). We assessed 217 patients and 171 of these patients were included in a prior publication (15).

*Statistical Analysis.* Two-sided unpaired *t*-test was used to compare the means of two groups if the variance was equal as determined by F-test. Mann-Whitney U-test was completed if there was unequal variance. Fisher's exact test was used to determine whether there was a difference in proportion of patients who are female sex between patients with hypochloremia and patients with normal serum chloride. Chi-squared test assessed whether there was a difference in PAH etiology between patients with hypochloremia and patients with normal serum chloride. One-way analysis of variance (ANOVA) with Tukey post-hoc analysis was performed to compare the means of three groups if there was equal variance as determined by the Brown-Forsythe test. Brown-Forsythe and Welch ANOVA with Dunnett post-hoc analysis were completed if there was unequal variance between groups. Simple linear regression evaluated whether there were differences in slopes and y-intercepts between the best fit lines of RA pressure *vs.* PVR and cardiac output *vs.* PVR in hypochloremic and normal chloride patients. Statistical significance was defined as $p<0.05$. Principal component analysis and hierarchical cluster analysis of the proteomic results were completed with MetaboAnalyst (55). Statistical analyses and graphing were performed with GraphPad Prism version 9. Data presented as mean ± standard deviation. Graphs show the mean and all individual values.

**Supplemental Table 1: Antibodies Used in Study.**

| Antigen | Company | Species | Catalogue Number | Dilution |
|---|---|---|---|---|
| **Western Blot** | | | | |
| AS160 | EMD Millipore | Rabbit | 07-741 | 1:250 |
| DJ-1 | Abcam | Rabbit | ab18257 | 1:250 |
| GFAT1 | Abcam | Rabbit | ab176775 | 1:100 |
| GLO1 | Thermo Fisher Scientific | Mouse | MA1-13029 | 1:100 |
| GLO2 | Thermo Fisher Scientific | Rabbit | PA5-93097 | 1:50 |
| GLUT1 | Abcam | Rabbit | ab115730 | 1:100 |
| GLUT4 | Cell Signaling Technology | Mouse | 2213S | 1:250 |
| Methylglyoxal | Cell Biolabs | Mouse | STA-011 | 1:100 |
| OGA | Abcam | Rabbit | ab124807 | 1:100 |
| OGT | Abcam | Rabbit | ab177941 | 1:250 |
| O-GlcNAc (RL2) | Thermo Fisher Scientific | Mouse | MA1-072 | 1:250 |
| Phospho-AS160 | Cell Signaling Technology | Rabbit | 8730S | 1:250 |
| WNK1 | Thermo Fisher Scientific | Rabbit | PA5-80229 | 1:100 |
| WNK2 | Abcam | Rabbit | ab192397 | 1:250 |
| Anti-mouse secondary | Li-COR | Goat | 926-32210<br>926-68070 | 1:5000 |
| Anti-rabbit secondary | Li-COR | Goat | 926-32211<br>926-68071 | 1:5000 |
| **Immunofluorescence** | | | | |
| GLUT1 | Abcam | Rabbit | ab115730 | 1:50 |
| GLUT4 | Abcam | Rabbit | ab654 | 1:50 |
| Succinylated WGA | Vector Laboratories | | FL-1021S | 1:50 |
| WNK1 | Thermo Fisher Scientific | Rabbit | PA5-80229 | 1:50 |
| Anti-rabbit secondary, Alexa Fluor plus 555 | Invitrogen | Goat | A32732 | 1:500 |
| Wheat germ agglutinin, Alexa Fluor 633 conjugate | Invitrogen | | W21404 | 1:50 |

**Supplemental Table 2: Clinical Characteristics of PAH Cohort.** Data shown as mean ± standard deviation. *P*-value determined by Fisher's exact test for proportion of patients who are female sex, chi-squared test for PAH etiology, and either *t*-test or Mann-Whitney U test if the data were not normally distributed. CTD: connective tissue disease; HIV: human immunodeficiency virus; NT-proBNP: N-terminal pro-brain natriuretic peptide; PAH: pulmonary arterial hypertension; PVOD: pulmonary veno-occlusive disease; RVFAC: right ventricular fractional area change; S': systolic velocity; TAPSE: tricuspid annular plane systolic excursion.

| **Characteristics** | **Hypochloremia (*n*=40)** | **Normal Chloride (*n*=177)** | ***P*-value** |
|---|---|---|---|
| Age, years | 58±14 | 53±17 | 0.06 |
| Female sex, *n* (%) | 30 (75%) | 127 (72%) | 0.85 |
| PAH etiology, *n* (%) | | | 0.03 |
| Idiopathic PAH | 14 (35%) | 33 (19%) | |
| Heritable PAH | 0 (0%) | 2 (1%) | |
| Drug-induced PAH | 0 (0%) | 13 (7%) | |
| CTD-associated PAH | 16 (40%) | 72 (41%) | |
| HIV-associated PAH | 1 (2.5%) | 2 (1%) | |
| Portopulmonary PAH | 7 (17.5%) | 26 (15%) | |
| Congenital heart disease | 0 (0%) | 25 (14%) | |
| PVOD | 0 (0%) | 2 (1%) | |
| Other | 2 (5%) | 2 (1%) | |
| Medications, *n* (%) | | | |
| Diuretics | 10 (25%) | 52 (29%) | 0.70 |
| Calcium channel blockers | 5 (12.5%) | 38 (21%) | 0.27 |
| Phosphodiesterase-5 inhibitors | 16 (40%) | 53 (30%) | 0.26 |
| Endothelin receptor antagonists | 7 (17.5%) | 16 (9%) | 0.15 |
| Prostacyclins | 5 (12.5%) | 13 (7%) | 0.34 |
| PAH-specific treatment naive | 0 (0%) | 1 (0.6%) | >0.99 |
| Laboratory | | | |
| Serum sodium, mmol/L | 135±4 (*n*=40) | 140±3 (*n*=166) | <0.0001 |
| Serum chloride, mmol/L | 98±3 (*n*=40) | 106±3 (*n*=168) | <0.0001 |
| Serum creatinine, mg/dL | 1.3±1.3 (*n*=40) | 1.0±0.7 (*n*=175) | 0.25 |
| Serum hemoglobin, g/dL | 12.4±2.1 (*n*=40) | 13.4±2.7 (*n*=172) | 0.03 |
| Serum NT-proBNP, pg/dL | 5095±10751 (*n*=37) | 2073±3588 (*n*=146) | 0.08 |
| Six-minute walk distance, m | 281±153 (*n*=21) | 350±118 (*n*=100) | 0.02 |
| Echocardiography | | | |
| RVFAC, % | 32±9 (*n*=24) | 34±11 (*n*=80) | 0.40 |
| TAPSE, cm | 1.8±0.6 (*n*=26) | 1.8±0.5 (*n*=96) | 0.55 |
| S', cm/s | 9.9±2.8 (*n*=20) | 10.7±2.5 (*n*=60) | 0.27 |
| Hemodynamics | | | |
| Heart rate, beats/min | 80±16 (*n*=31) | 77±15 (*n*=151) | 0.24 |
| Mean right atrial pressure, mmHg | 10±6 (*n*=39) | 7±5 (*n*=173) | 0.002 |
| Mean pulmonary artery pressure, mmHg | 43±10 (*n*=40) | 45±14 (*n*=176) | 0.71 |
| Pulmonary capillary wedge pressure, mmHg | 10±3 (*n*=40) | 9±3 (*n*=177) | 0.09 |
| Cardiac output, L/min | 5.0±2.0 (*n*=24) | 4.9±1.9 (*n*=131) | 0.74 |
| Cardiac index, L/min/m$^2$ | 2.5±0.9 (*n*=24) | 2.6±1.0 (*n*=129) | 0.65 |
| Pulmonary vascular resistance, Wood units | 8.6±4.4 (*n*=38) | 9.1±5.7 (*n*=165) | 0.62 |
| Pulmonary arterial compliance, mL/mmHg | 1.7±1.0 (*n*=30) | 1.7±1.2 (*n*=146) | 0.97 |

**Supplemental Table 3: Transcript Expression of WNK1, GLUT1, GLUT4, and AS160 in Human LV and RV Cardiomyocytes.** Data obtained from the Human Cell Atlas (36), which contains single-cell and single-nucleus RNA sequencing data from donor hearts. Five ventricular cardiomyocyte populations (vCM1-5) are identified in the Human Cell Atlas and the RV is more enriched with group 2 cardiomyocytes (vCM2_RV) than the LV (39.91 *vs.* 9.12%). WNK1, GLUT4, and AS160 are expressed slightly higher in vCM2_RV than vCM2_LV.

| Gene | vCM1_LV | vCM1_RV | vCM2_LV | vCM2_RV | vCM3_LV | vCM3_RV | vCM4_LV | vCM4_RV | vCM5_LV | vCM5_RV |
|---|---|---|---|---|---|---|---|---|---|---|
| **WNK1** | 2.682 | 2.814 | 2.835 | 2.964 | 2.588 | 2.38 | 2.195 | 2.487 | 3.12 | 3.072 |
| **GLUT1/SLC2A1** | 0.059 | 0.057 | 0.068 | 0.05 | 0.066 | 0.055 | 0.114 | 0.129 | 0.011 | 0.073 |
| **GLUT4/SLC2A4** | 0.776 | 0.798 | 0.697 | 0.861 | 0.542 | 0.586 | 0.918 | 1.061 | 0.705 | 0.843 |
| **AS160/TBC1D4** | 4.898 | 5.581 | 4.592 | 5.182 | 4.583 | 5.688 | 3.822 | 4.932 | 4.692 | 5.132 |

**Supplemental Figure 1: Complete Western Blot Images.** Complete Western blot images for all proteins assessed in this study.

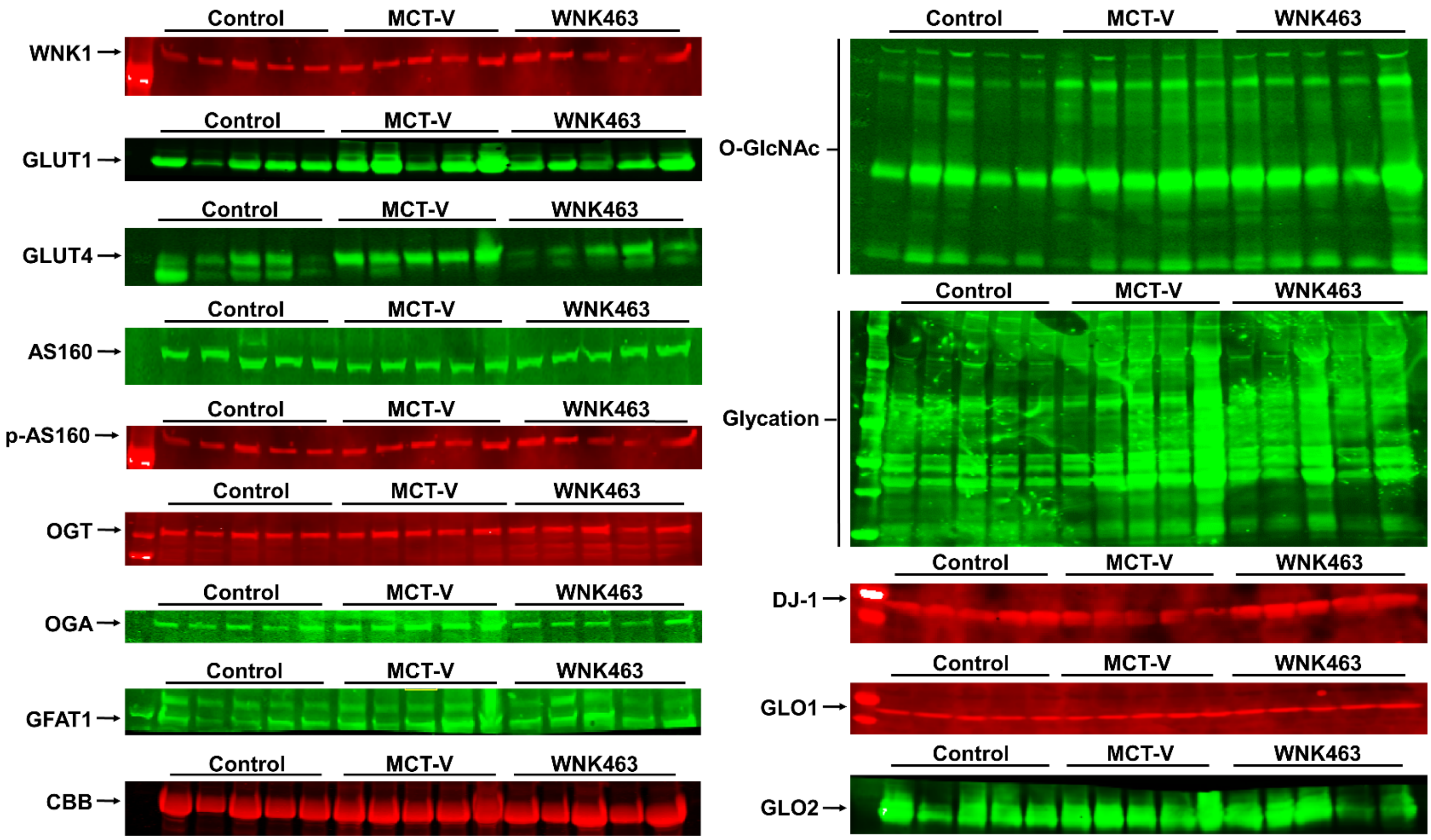

**Supplemental Figure 2: Representative Echocardiogram Images.** Representative echocardiogram images and measurements from control, MCT-V, and WNK463.

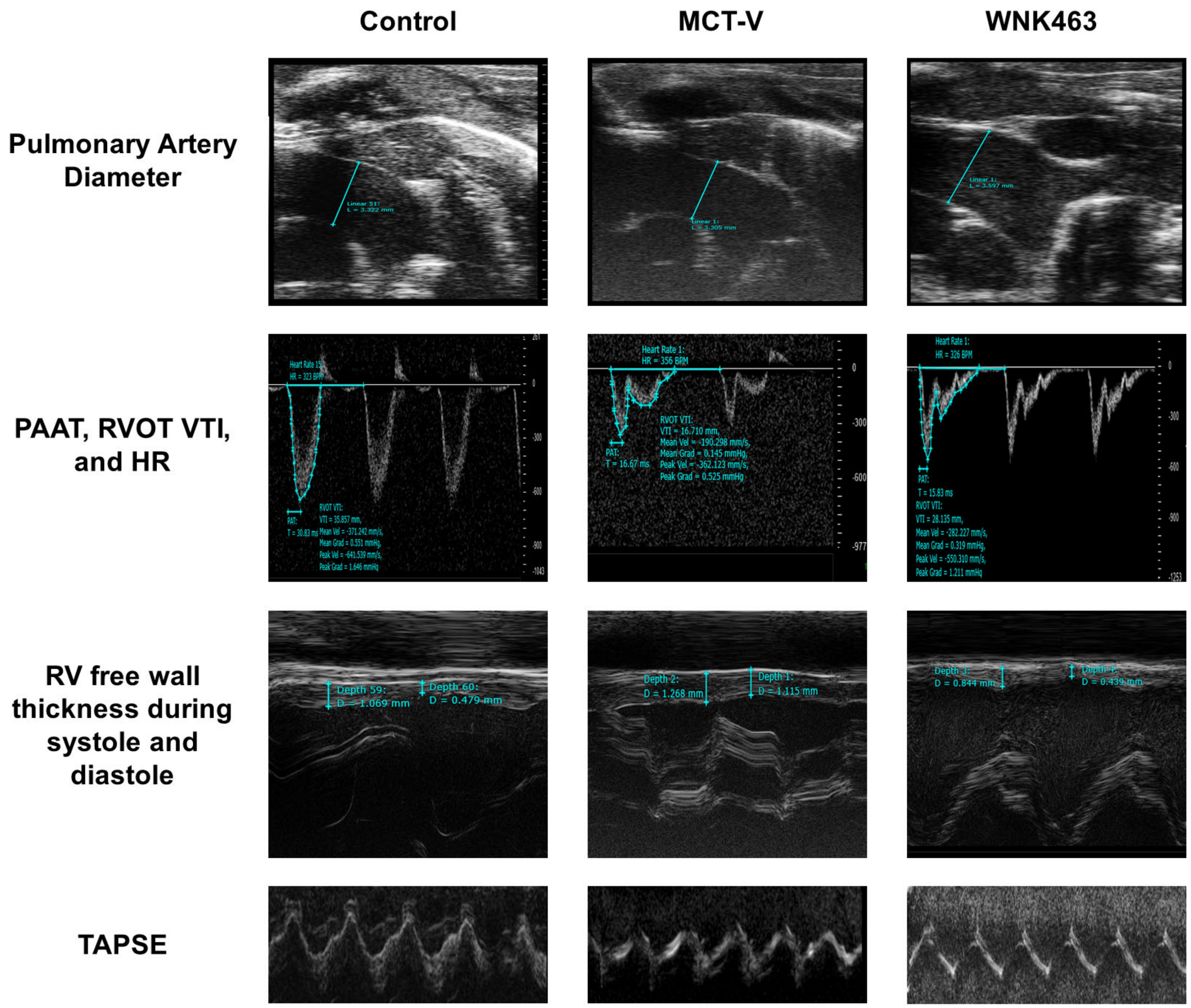

**Supplemental Figure 3: Representative Pressure-Volume Loops.** Representative PV loops from control, MCT-V, and WNK463. End-Systolic Elastance (Ees) line (red) shown.

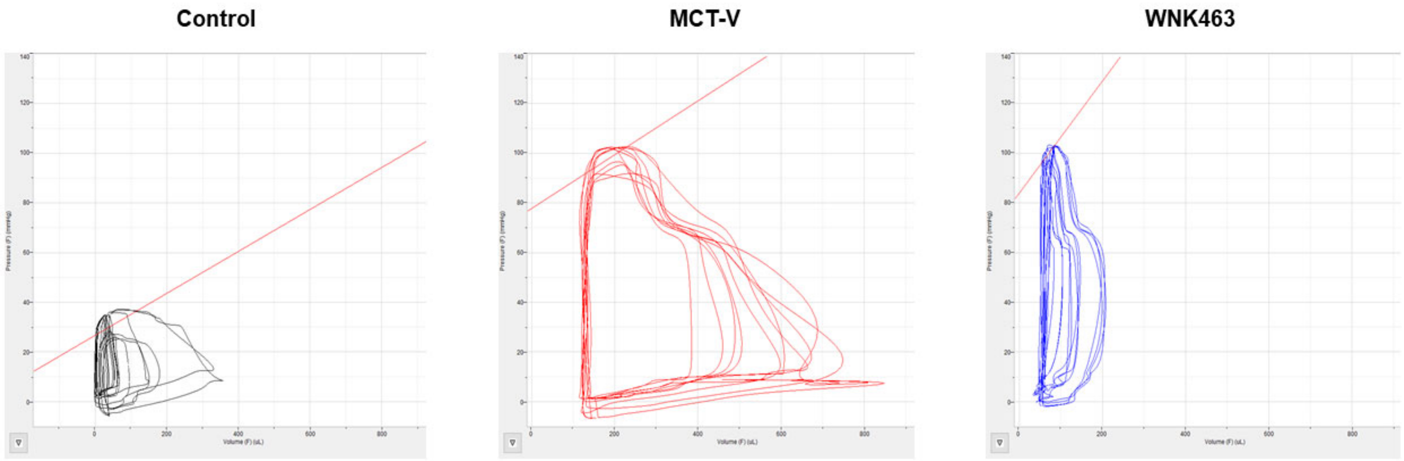

**Supplemental Figure 4: Integration of RV Mitochondrial/Peroxisomal Proteomics and RV Metabolomics Data Highlighting Fatty Acid Oxidation as the Most Altered Metabolic Pathway in MCT PAH RV.** The three most dysregulated pathways identified are fatty acid degradation, the TCA cycle, and pyruvate metabolism.

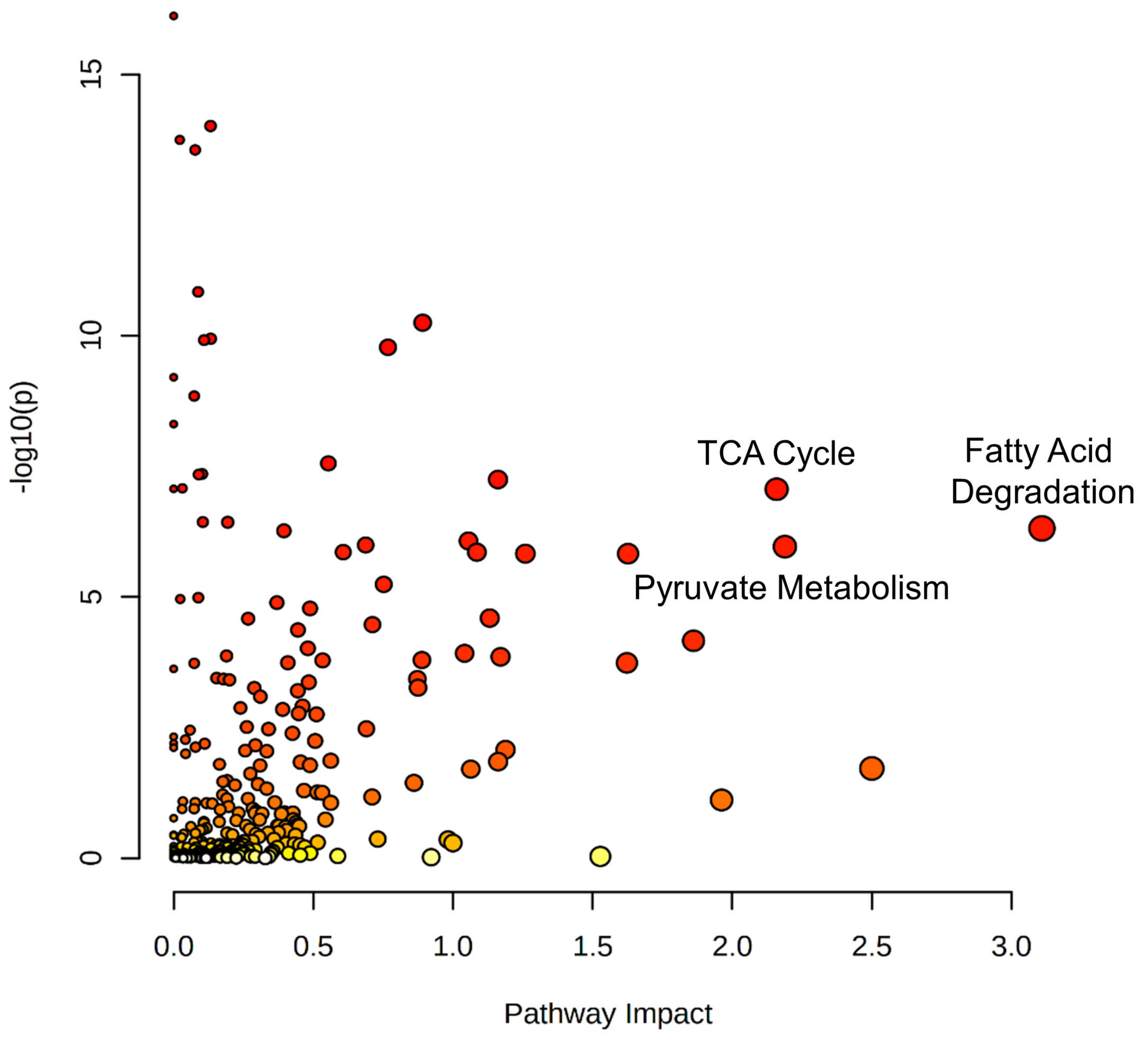

**Supplemental Figure 5: Multiple Peroxisomal Proteins are Elevated in MCT-V and WNK463 RVs.**
Hierarchical cluster analysis of peroxisomal proteins in proteomics experiments.

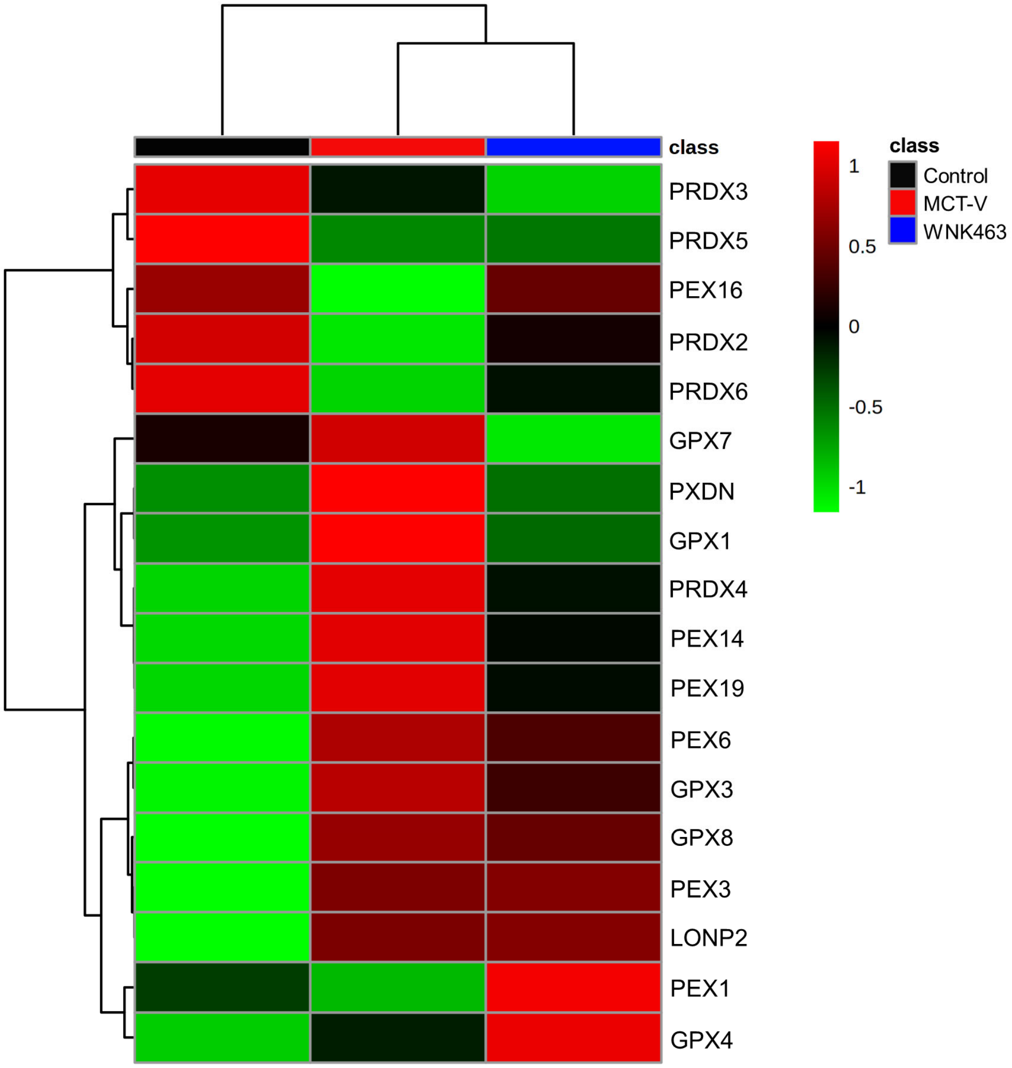

**Supplemental Figure 6: WNK2 Expression is detected in brain but RV extracts.** Representative Western blots from RV and brain extracts.

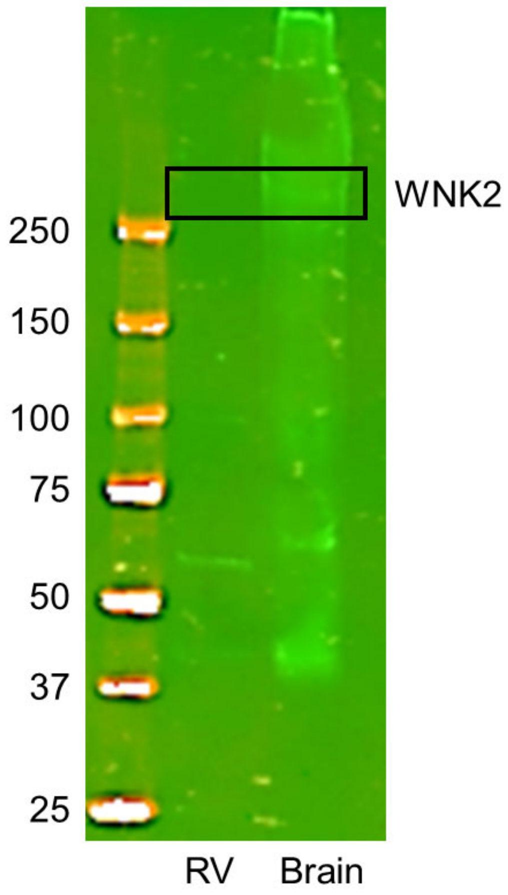

**Supplemental Figure 7: Proteomic Assessment of Mitochondrial Ribosomal and Membrane Proteins.** As shown by hierarchical cluster analysis of mitochondrial/peroxisomal proteomics studies, most mitochondrial ribosomal and membrane proteins are elevated in MCT-V and WNK463 RVs.

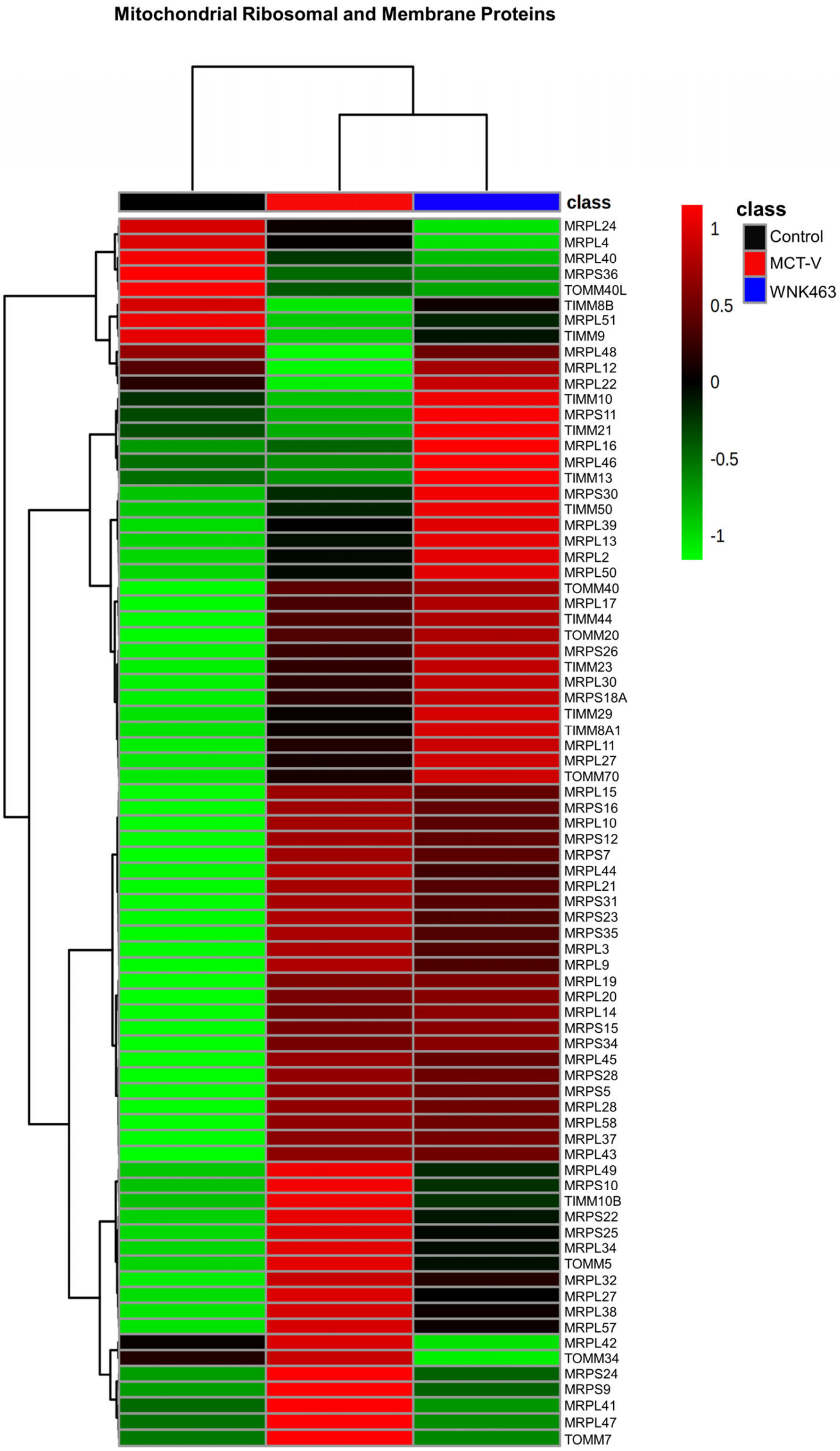

**Supplemental Figure 8: WNK463 Mitigates the Development of RV Fibrosis.** As shown in representative images of RV tissue in (**A**) and quantified in (**B**), WNK463 reduces the amount of fibrosis (arrows) in the RV (Control: 0.4±0.2%, MCT-V: 2.2±1.3%, WNK463: 1.0±0.7%, $p$=0.04 between MCT-V and WNK463, $n$=11-12 total areas assessed from 3 RVs per group). Scale bar 50 μm. *$p$<0.05 and **$p$<0.01 by Brown-Forsythe and Welch ANOVA with Dunnett's multiple comparisons test.

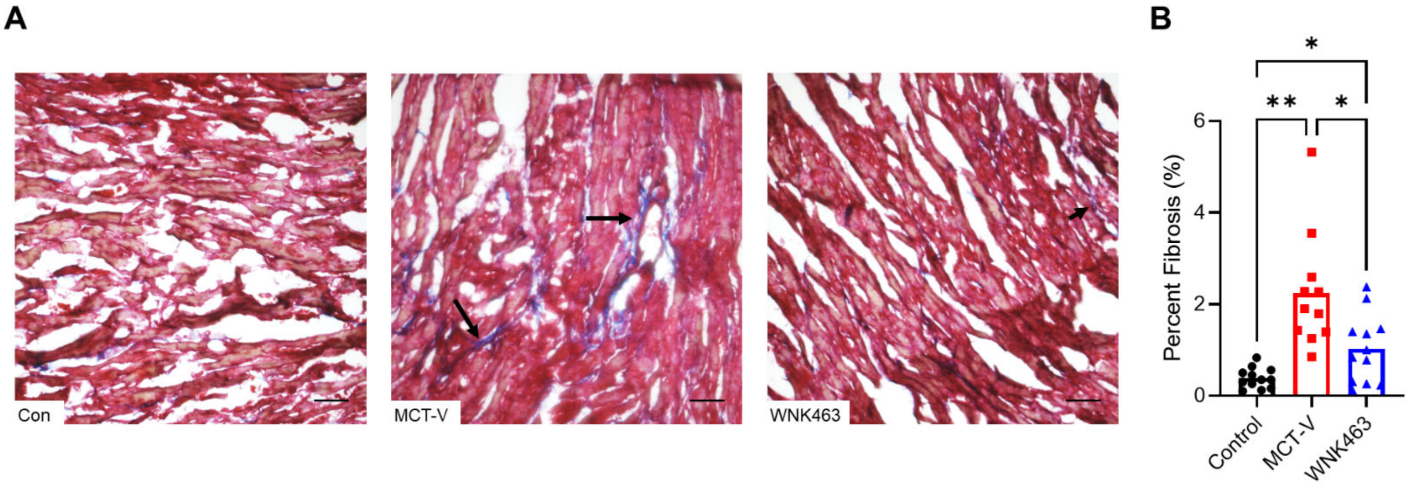
